# Medulloblastoma oncogene aberrations are not involved in tumor initiation, but essential for disease progression and therapy resistance

**DOI:** 10.1101/2024.02.09.579690

**Authors:** Konstantin Okonechnikov, Piyush Joshi, Verena Körber, Anne Rademacher, Michele Bortolomeazzi, Jan-Philipp Mallm, Patricia Benites Goncalves da Silva, Britta Statz, Mari Sepp, Ioannis Sarropoulos, Tetsuya Yamada-Saito, Jan Vaillant, Andrea Wittmann, Kathrin Schramm, Mirjam Blattner-Johnson, Petra Fiesel, Barbara Jones, Till Milde, Kristian Pajtler, Cornelis M. van Tilburg, Olaf Witt, Konrad Bochennek, Katharina Johanna Weber, Lisa Nonnenmacher, Christian Reimann, Ulrich Schüller, Martin Mynarek, Stefan Rutkowski, David T.W. Jones, Andrey Korshunov, Karsten Rippe, Frank Westermann, Supat Thongjuea, Thomas Höfer, Henrik Kaessmann, Lena M. Kutscher, Stefan M. Pfister

## Abstract

Despite recent advances in understanding disease biology, treatment of Group 3/4 medulloblastoma remains a therapeutic challenge in pediatric neuro-oncology. Bulk-omics approaches have identified considerable intertumoral heterogeneity in Group 3/4 medulloblastoma, including the presence of clear single-gene oncogenic drivers in only a subset of cases, whereas in the majority of cases, large-scale copy-number aberrations prevail. However, intratumoral heterogeneity, the role of oncogene aberrations, and broad CNVs in tumor evolution and treatment resistance remain poorly understood. To dissect this interplay, we used single-cell technologies (snRNA-seq, snATAC-seq, spatial transcriptomics) on a cohort of Group 3/4 medulloblastoma with known alterations in the oncogenes *MYC, MYCN*, and *PRDM6*. We show that large-scale chromosomal aberrations are early tumor initiating events, while the single-gene oncogenic events arise late and are typically sub-clonal, but *MYC* can become clonal upon disease progression to drive further tumor development and therapy resistance. We identify that the subclones are mostly interspersed across tumor tissue using spatial transcriptomics, but clear segregation is also present. Using a population genetics model, we estimate medulloblastoma initiation in the cerebellar unipolar brush cell-lineage starting from the first gestational trimester. Our findings demonstrate how single-cell technologies can be applied for early detection and diagnosis of this fatal disease.

## Main

Intratumoral heterogeneity, a hallmark of cancer, refers to diverse molecular and functional cell populations within a single tumor^1^. Intratumoral heterogeneity is driven by genetic mutations, transcriptomic/ epigenomic plasticity, and reprogramming of microenvironment^2^. The malignant childhood tumor medulloblastoma is heterogeneous^3^, especially in the Groups 3 and 4 . This heterogeneity makes effective treatment of these tumors difficult and contributes to overall low survival rates^4^. Advanced DNA methylation profiling has classified Group 3/4 tumors into 8 distinct molecular subgroups^5^. In addition, single-cell transcriptomic profiling has unveiled the intricate regulatory activity of transcription factors and signaling pathways that orchestrate cellular diversity^6,7^. Despite these advances, the role of oncogenes in shaping intratumor heterogeneity remains unknown.

A minority of Group 3/4 medulloblastoma tumors harbor single-gene oncogenic drivers, including *MYC*^*8*^ and *MYCN*^*9*^ amplifications as well as *PRDM6* overexpression due to enhancer hijacking via a tandem duplication of the adjacent *SNCAIP* gene^3^. In contrast, the majority of these tumors display recurrent, large-scale copy number changes^3,10,11^, including chromosome 8 and 11 loss, and chromosome 7 gain and isochromosome 17q. A fundamental question of which genetic events initiate and drive these tumors remains unanswered. Using single-cell multi-omic and spatial transcriptomic approaches, we determined the interplay between large-scale copy number variants and single-gene somatic events in driving medulloblastoma heterogeneity and evolution.

### Driver oncogenic events in primary tumors are typically subclonal with distinct molecular properties

To understand the clonal genetic events in tumor initiation, evolution, and progression, we molecularly profiled a specific tumor cohort with a known amplification of *MYCN* or *MYC* or overexpression of *PRDM6* (n=16 primary, n=4 relapses, **Fig. 1a, Extended Data Table 1**, dataset published in Joshi *et al bioRxiv*). In larger datasets, amplification or activation of these oncogenes is present in approximately 30% of Group 3/4 medulloblastoma cases^3^, whereas ∼70% of cases lack a single-gene somatic event. The presence of focal *MYC*/*MYCN* amplifications or the *SNCAIP* tandem duplication was verified using bulk molecular profiles or fluorescence in situ hybridization (FISH) for each sample in this cohort (**Extended Data Table 1**). We analyzed single-nucleus profiles of our target cohort, examining both RNA-sequencing (snRNA-seq, n=20) and the matched ATAC-sequencing (snATAC-seq, n=16) in the same nuclei.

**Figure 1.**
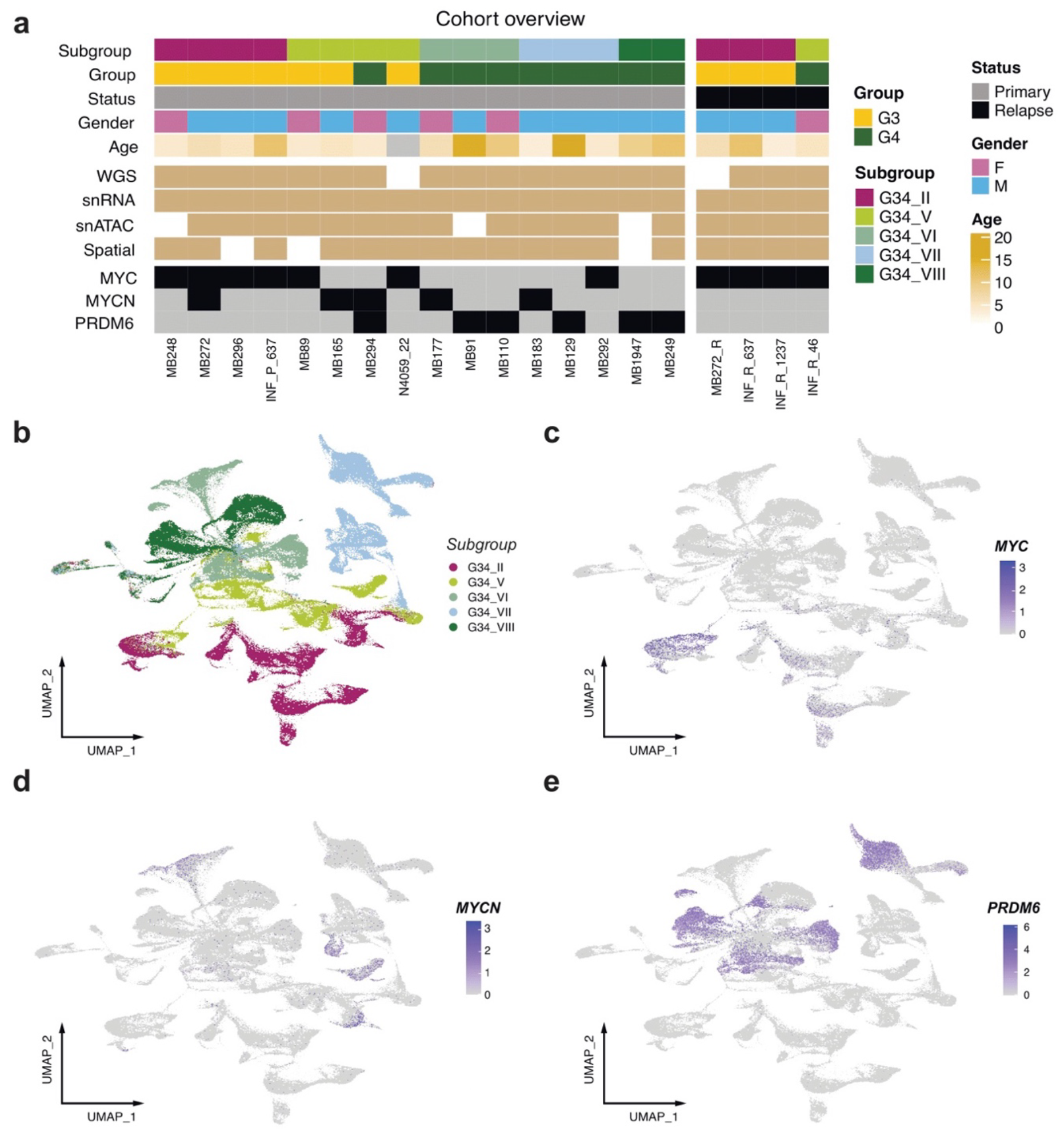
Single-nucleus transcriptional profiling of 16 oncogene-driver Group 3/4 medulloblastoma primary tumor samples. a) Overview of target cohort with annotation. Two primary-relapse pairs (MB272/R, INF_P/R_637) are from the same patients. b) UMAP of snRNA-seq merged dataset, medulloblastoma subgroups annotated. Feature plots showing c) *MYC*, d) *MYCN*, e) *PRDM6* expression within UMAP of merged snRNA-seq dataset.

Uniform manifold approximation and projection (UMAP) visualization of the snRNA-seq data from primary tumors showed Group 3/4 subgroup-specific clusters without batch effect adjustments (**Figure 1b**); mixed normal cell types arising from different samples clustered together as expected (**Extended Data Fig. 1a**). Expression of the oncogenes *MYC, MYCN* and *PRDM6* demonstrated clear sample specificity (**Fig. 1c-e, Extended Data Fig. 1b**). In addition, expression of known marker genes delineated non-tumor cell types, including *PTPRC* (microglia), *IGFBP7* (meningeal), and *AQP4* (astroglia) (**Extended Data Fig. 1c-e**). This separation of structure was recapitulated with snATAC-seq, as visualized via UMAP (**Extended Data Fig. 1f,g**). For samples with multi-omic data, non-tumor cells in snATAC-seq were labeled based on their associated non-tumor clusters from the snRNA-seq data (**Extended Data Fig. 1f**).

To investigate the clonal heterogeneity of *MYCN*-amplified tumor samples (subgroups V/VII), we adapted the inferCNV^12^ approach (see Methods) to infer copy number variation (CNV) profile of cell-clusters, using both snRNA-seq and snATAC-seq data. We identified common CNV changes across all tumor cells, which were also verified with bulk data (**Extended Data Table 1**); however, in most cases, we also observed clusters with discordant CNVs, which we labelled as subclones. For example, in all *MYCN-*amplified tumors (n=4), we identified two distinct subclones: with (C1) and without (C2) *MYCN* amplification, respectively (**Fig. 2a, Extended Data Fig. 2a**). Remarkably, in all cases, reconstruction of the putative phylogenetic trees showed that *MYCN* amplification was not the initiating event for the tumor. Instead, large-scale CNVs, such as loss of chromosomes 8 and 10 or gain of chromosomes 7 or 17q, were already present in the presumptive founder clone (C0) (**Fig. 2b**). Moreover, additional unique CNVs were found only within *MYCN*-and non-*MYCN*-amplified subclones.

**Figure 2.**
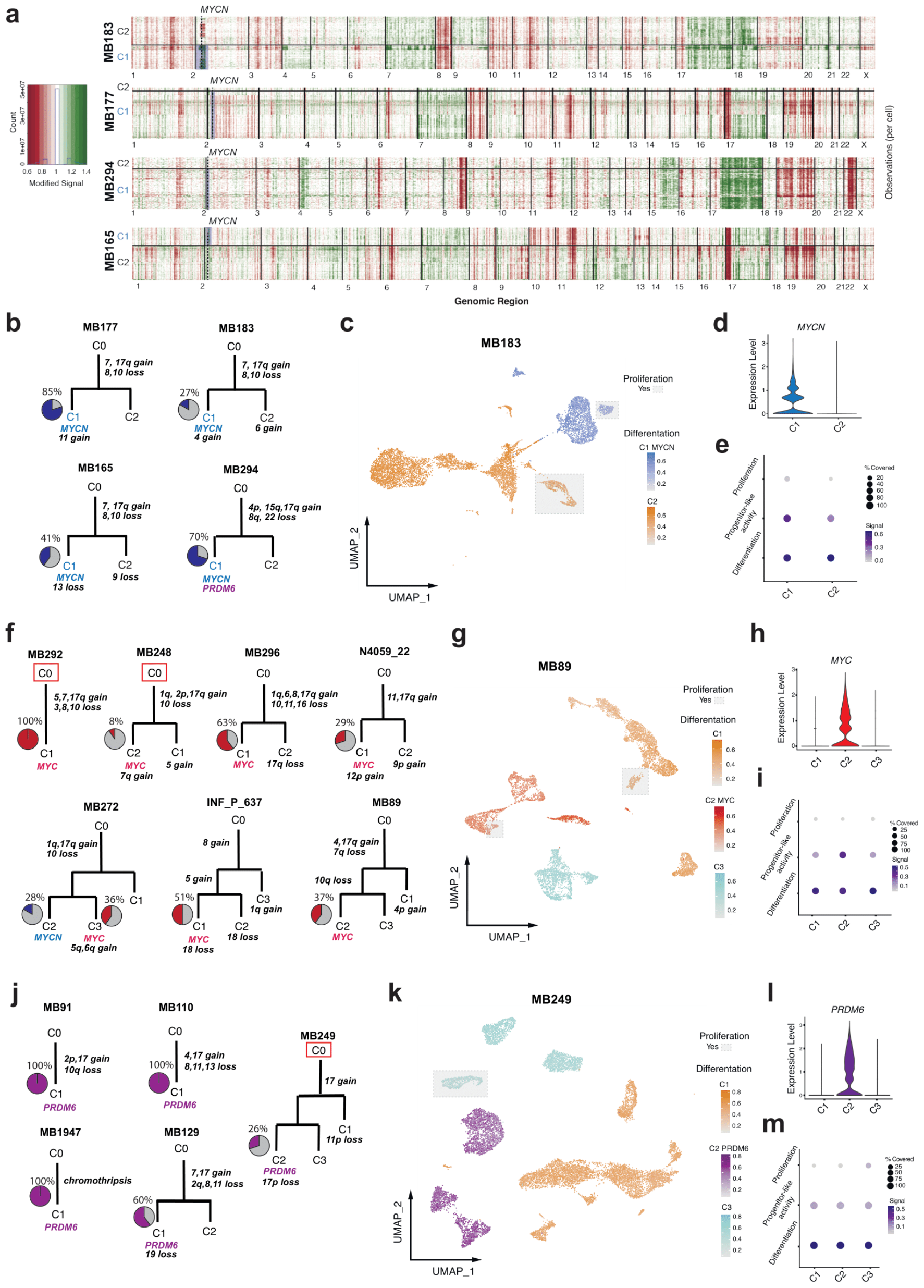
Clonal proliferation and differentiation gradients are independent of oncogene expression. a) Copy number of snATAC-seq data from *MYCN* samples. Red, chromosome loss. Green, chromosome gain. b) Somatic phylogeny trees for *MYCN* samples. Blue, proportion of *MYCN*-expressing cells. c) snRNA-seq UMAP of single *MYCN* sample MB183. Gray boxes, proliferating cells. Blue, differentiation signal enrichment in *MYCN*-expressing C1 clone. Orange, differentiation signal enrichment in C2 clone. d) *MYCN* expression in C1 and C2 clones. e) Per cell gene set variance analysis (GSVA) enrichments of proliferating, progenitor-like and differentiation in single sample shown in c. f) Somatic phylogeny trees for *MYC* samples. Red, proportion of *MYC*-expressing cells. Red square, cases MB292 and MB248 with somatic mutations in C0. g) snRNA-seq UMAP of single MYC sample MB89. Gray boxes, proliferating cells. Red, differentiation signal enrichment in *MYC*-expressing C2 clone. Orange, differentiation signal enrichment in C1 clone. Aquamarine, differentiation signal enrichment in C3 clone. h) *MYC* expression in C1, C2, and C3 clones. i) Per cell GSVA enrichments of proliferating, progenitor-like and differentiation in single sample shown in g. j) Somatic phylogeny trees for *PRDM6* samples. Purple, proportion of *PRDM6*-expressing cells. Red square, case MB249 with somatic mutations outside of CNV regions. k) snRNA-seq UMAP of single *PRDM6* sample MB249. Gray boxes, proliferating cells. Purple, differentiation signal enrichment in *PRDM6*-expressing C2 clone. Orange, differentiation signal enrichment in C1 clone. Aquamarine, differentiation signal enrichment in C3 clone. l) *PRDM6* expression in C1, C2, and C3 clones. m) Per cell GSVA enrichments of proliferating, progenitor-like and differentiation in single sample shown in k.

Next, we examined the differentiation, proliferation, and aggressive progenitor-like activity states of individual cells within each subclone using snRNA-seq expression of established reference gene lists for these defined medulloblastoma cell states^6^. We identified that both *MYCN*-amplified and non-amplified subclones maintained separate proliferating and differentiated compartments (**Fig. 2c,d, Extended Data Fig. 2b)**. The *MYCN* subclone was also uniquely enriched with a progenitor-like gene expression signature (**Fig. 2e**). As cells differentiated, expression of the oncogene itself showed lower expression within the *MYCN-*amplified subclone (Pearson cor. = -0.23, p < 2.2e-16, **Extended Data Fig. 2c**). Similar differentiation levels among subclones and a slight bias towards progenitor activity in *MYCN*-amplified subclones was observed in all four *MYCN*-amplified tumors **(Extended Data Fig. 2d**).

Performing single-cell CNV analyses on *MYC*-amplified tumor samples (subgroups II/V), we identified a subclonal *MYC* amplification in 6 out of 7 samples (**Fig. 2f, Extended Data Fig. 2e**). Similar to *MYCN*-amplified tumors, the common and likely initiating events in the founder clone (C0) were large-scale chromosome 10 loss and/or chromosome 17q gain, with subclonal *MYC-*amplification occurring later during tumor evolution. Remarkably, the clonal-structure of *MYC-*amplified tumors was more complex (N=3/7 cases), with the formation of three or more unique subclones (**Fig. 2f** bottom). Typically, *MYC*-amplified subclones had their own proliferating and differentiating compartments (**Fig. 2g**,**h; Extended Data Fig. 2f**). Similar to the *MYCN-*amplified clones, only *MYC-*amplified clones demonstrated strong enrichment of progenitor-like activity compared to non-*MYC-*amplified compartments (**Fig. 2i**). Differentially expressed genes specific to *MYC*-amplified subclones were enriched in known *MYC* target genes^13^ (p<1.11e-16, **Extended Data Table 2**) and *MYC* expression decreased as cells differentiated (Pearson cor. = -0.18, p < 2.2e-16, **Extended Data Fig.2g**). Across six samples with subclonal *MYC* amplifications, the differentiation level was similar among subclones; however, progenitor-like activity was significantly enriched in *MYC-*amplified subclones (**Extended Data Fig.2h**)

Lastly, we examined tumors with enhanced *PRDM6* expression (subgroups VII / VIII), where *SNCAIP* gene duplication leads to aberrant activation of *PRDM6* via enhancer hijacking^3^. In our cohort, we identified 2 out of 5 samples where *PRDM6* overexpression was subclonal (**Fig. 2j, Extended Data Fig. 3a**). Chromosome 17q gain was the most frequent CNV within the founder clone (C0). In contrast to *MYC* and *MYCN* clones, we could not identify a distinct proliferating compartment in *PRDM6*-specific clones (**Fig. 2k,l**). Instead, we only found an overall small proportion of cells (< 5%) with the proliferation gene signature in *PRDM6* subclones (**Extended Data Fig. 3b)**. We also did not identify enriched progenitor-like activity in the *PRDM6* subclones (**Fig. 2m; Extended Data Fig. 3c**), except in one specific case where we detected a *MYCN-*amplified subclone that additionally harbored a *SNCAIP* duplication with associated *PRDM6* overexpression (**Fig. 2b**, bottom right).

Despite the low mutational burden in medulloblastoma^3^, we also inspected the somatic single-nucleotide variants (SNVs) to confirm the tumor phylogeny composition predicted by snRNA-and snATAC-seq data. Using whole-genome sequencing (WGS) data, we examined mutations in CNV regions specific to the founder clone (C0) or not lying within CNV. In 9/12 samples, we did not identify the presence of drivers or co-mutations (exception: 2 *MYC*-amplified cases, **Fig 2f** and 1 PRDM6 case, **Fig 2j**).

Collectively, these findings nominate large-scale CNVs as likely tumor-initiating events in group 3/4 medulloblastoma with focal oncogene aberrations only occurring during tumor evolution.

### Clonal mutation densities suggest Group 3/4 medulloblastoma onset occurs from first trimester until first year of life

To investigate clonal evolution during the initiation of Group 3/4 medulloblastomas, we analyzed WGS data from the medulloblastoma ICGC cohort^3^ (**Extended Data Table 3**). Somatic tissues accumulate SNVs continuously over time^14-16^, and hence SNV density in the tumor cell of origin (in population genetics “most recent common ancestor”, MRCA) can be interpreted as a measure for the patient’s age at tumor initiation^17,18^. To time the developmental origin of medulloblastoma with this approach, we quantified clonal SNV densities from the allele frequency distribution of somatic variants in 183 primary medulloblastomas of all subgroups (**Extended Data Fig. 4a and b**; comprising 109 Group 3/4 medulloblastomas, 21 infant Sonic Hedgehog (SHH)-medulloblastomas, 36 childhood/adulthood SHH-medulloblastomas and 17 WNT-medulloblastomas). Overall, the clonal SNV densities across subgroups recapitulated the age-incidence distribution of the disease, with infant SHH-medulloblastoma having lowest densities (0.02±0.01 SNVs/Mb), followed by Group 3/4 medulloblastoma (0.1±0.08 SNVs/Mb), WNT-medulloblastoma (0.28±0.47 SNVs/Mb) and adult SHH-medulloblastoma (0.41±0.45 SNVs/Mb; **Extended Data Fig. 4c**). Clonal SNV densities were also correlated with age at diagnosis (Spearman’s rho = 0.73, p < 2.2e-16; **Extended Data Fig. 4d**), collectively supporting our approach to infer the evolutionary dynamics at medulloblastoma onset from somatic SNVs.

To estimate age of tumor initiation in Group 3/4 medulloblastomas, we analyzed 109 tumor samples of this subgroup in more detail (**Fig. 3a**). As with the entire cohort, clonal SNV densities were likewise correlated with the age at diagnosis among Group 3/4 medulloblastoma (Spearman’s rho = 0.52, p = 1.402e-08; **Extended Data Fig. 4e**). However, contrary to the clear temporal order in tumor initiation of the major medulloblastoma groups, clonal SNV densities were statistically indistinguishable between Group 3/4 medulloblastoma subgroups I-VIII (Wilcoxon rank sum test, all adjusted p values > 0.05), indicating that growth of the final tumor mass commences around the same developmental time window in all Group 3/4 medulloblastoma subgroups (**Figure 3b**).

**Figure 3.**
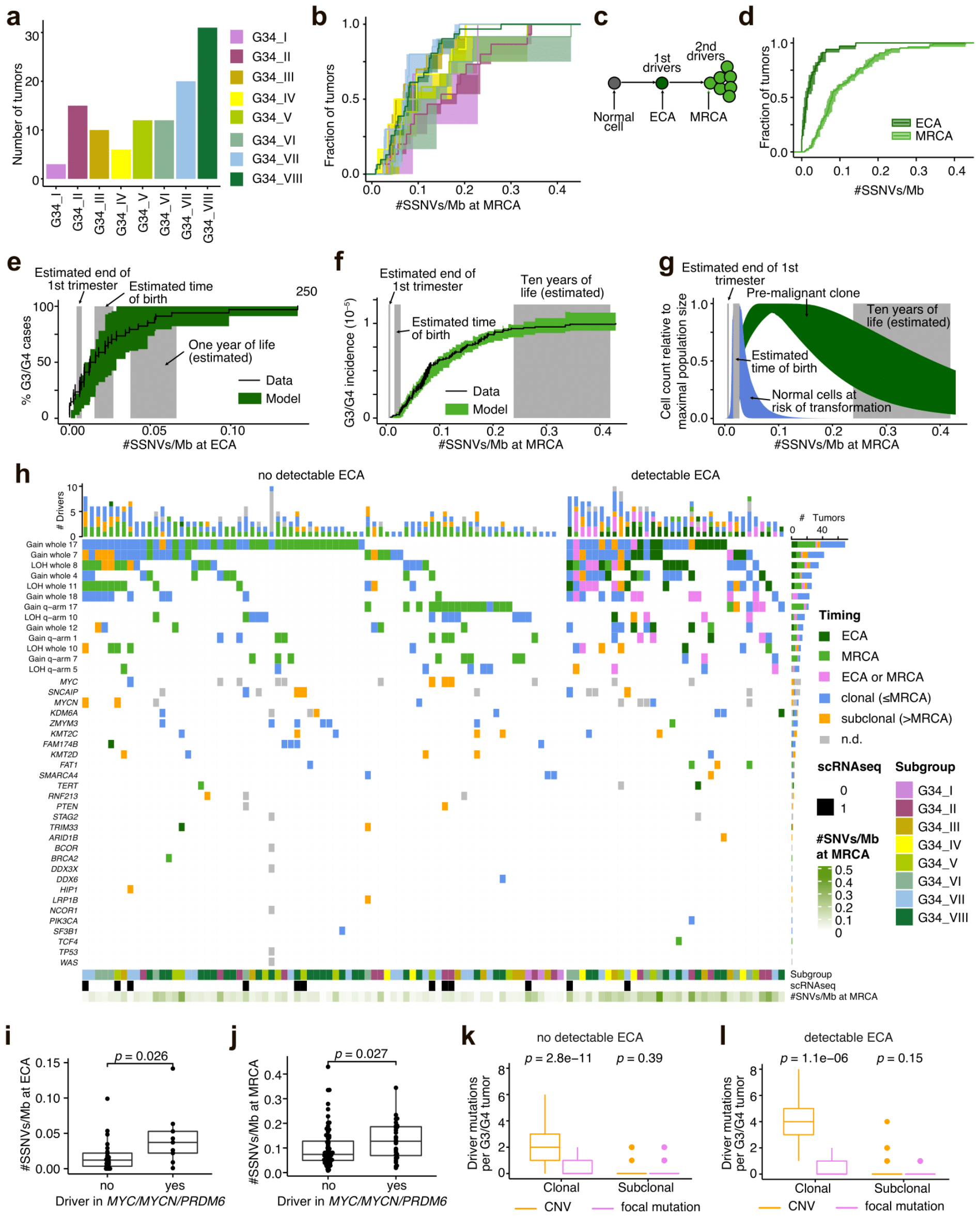
Development of Group 3/4 medulloblastoma somatic mutation profiles and association with cells-of-origin. a) Overview of Group 3/4 medulloblastoma (MB G34) analyzed by bulk WGS and stratified by subgroup. b) SNV densities at MRCA stratified by subgroup (MB G34 I, n = 3, MB G34 II, n = 15, MB G34 III, n = 10, MB G34 IV, n = 6, MB G34 V, n = 12, MB G34 VI, n = 12, MB G34 VII n = 20, MB G34 VIII, n = 31). Shown are mean and 95% CI (estimated by bootstrapping the genomic segments 1,000 times; see Methods). c) Early medulloblastoma evolution. Initial acquisition in an early common ancestor (ECA) leads to a pre-malignant lesion, which transforms into a malignant clone upon acquisition of additional driver genes in the tumor’s most recent common ancestor (MRCA). d) SNV densities at ECA and MRCA for Group 3/4 medulloblastoma (n = 109). Mean and 95% CI, estimated by bootstrapping the genomic segments 1,000 times. e) Model fit to SNV densities at ECA. Line, mean and 95% CI (estimated by bootstrapping the genomic segments 1,000 times; see Methods) of the measured SNV densities; shaded area, 95% credible interval. 95% credible interval of the estimated time of birth is shown in grey. Second x-axis (estimated weeks p.c.) was computed as detailed in Methods. f) Model fit to SNV densities at MRCA. Line, mean and 95% CI of the measured SNV densities; shaded area, 95% credible interval of the estimated time of birth. Second x-axis, estimated weeks post conception (p.c.). g) Modeled tissue of origin (95% credible interval; dark grey), pre-malignant clone (95% credible interval; green) and estimated time of birth (95% credible interval). Cell counts relative to the maximal population size. Second x-axis, estimated weeks p.c. h) Genetic aberration spectrum in Group 3/4 medulloblastoma, stratified by timing (ECA, CNV could be uniquely timed to ECA; MRCA, CNV could be uniquely timed to MRCA; ECA or MRCA; SNV density at CNV in agreement with ECA and MRCA; clonal, CNV or small mutation was clonal without more detailed time information; subclonal, CNV or small mutation was subclonal; n.d., no data). Subclonality of *MYC* amplification, *MYCN* amplification and duplication of SNCAIP were integrated from the single-cell data. Note that mutations in *SNCAIP* lead to *PRDM6* overexpression. i) SNV density at ECA in Group 3/4 medulloblastoma with aberrations in *MYC/MYCN* or *PRDM6*. P value, unpaired Wilcoxon rank sum test (n = 81 without and n = 28 with aberration). j) SNV density at MRCA in Group 3/4 medulloblastoma with *MYC/MYCN* or *PRDM6* aberrations. p value, Wilcoxon rank sum test (n = 81 without and n = 28 with aberration).k) Number of clonal and subclonal CNV drivers and small mutations in known tumor-related genes among Group 3/4 medulloblastoma without detectable ECA. p values, paired Wilcoxon rank sum test (n = 75). l) Number of clonal and subclonal CNV drivers and small mutations in known tumor-related genes among Group 3/4 medulloblastoma with detectable ECA. p values, paired Wilcoxon rank sum test (n = 35).

To refine our analysis, we timed the acquisition of CNVs (clonal copy number gains or loss of heterozygosity, LOH) relative to the tumor’s MRCA. To this end, we compared densities of clonal SNVs acquired prior to a chromosomal gain, and hence present on multiple copies of a chromosomal region, to the density of clonal SNVs overall (Methods). Similar to neuroblastomas^18^ and other tumor entities^17^, 34 out of 109 Group 3/4 medulloblastomas showed evidence for having acquired at least one copy number gain in an early common ancestor (ECA), antecedent to the tumor’s MRCA (**Fig. 3c,d**). Hence, our data suggest that at least some CNVs in Group 3/4 medulloblastoma arise prior to the onset of tumor growth, in line with multiple rounds of mutation and selection at tumor initiation.

To date these events in actual time, we calibrated a population-genetics model of mutation and selection during tumor initiation^18^ with the measured SNV densities at ECA and MRCA, along with the patient age at diagnosis (see Methods for details). Briefly, the model assumes that medulloblastoma initiation is driven by clonal selection for two consecutive drivers, associated, respectively, with ECA and MRCA, in the transient cell population of (differentiating) unipolar brush progenitor cells from the rhombic lip^19-21^ (**Extended Data Fig. 4f**). Upon malignant transformation of the tumor’s MRCA, we assume exponential growth until a tumor size of 10^9^ cells (corresponding to a few cubic centimeters) is reached at the age of diagnosis (see Methods for details^22^). By fitting this model to the clonal SNV densities measured in Group 3/4 medulloblastomas, we inferred that the first oncogenic event (*i.e*., the ECA) occurs within the first gestational trimester in 24% of the cases, during late gestation in around 35% of cases and within the first year of life in 20% of cases (**Extended Data Fig. 4g, Fig. 3e**). The onset of tumor growth from its MRCA is placed considerably later, within the first decade of life (**Fig. 3f**), suggesting a long latency phase between pre-malignancy and the detection of a symptomatic tumor. Overall, the inferred dynamics of tumor initiation are consistent with a tumor origin in (differentiating) unipolar brush progenitor cells^19,23^, sustaining a pre-malignant clone that outlives the cell state of origin for several years (**Fig. 3g**).

### Large-scale copy number variation drives early tumor growth

To gain mechanistic insight into Group 3/4 medulloblastoma initiation, we asked whether particular mutations occur predominantly early or late. To address this question, we first focused on CNVs that were found more frequently than expected by chance, and hence are likely drivers of malignancy. Combining the enrichment results obtained in our cohort (**Extended Data Fig. 4h** and Methods) with published data^24^, we classified gains of Chromosome 1q, 4, 7, 12, 17/17q and 18, as well as LOH on Chromosome 5q, 8, 10/10q, 11 and 17p as putative drivers of Group 3/4 medulloblastoma initiation. Except for four cases, in which no ECA was identifiable, all Group 3/4 medulloblastomas harbored at least one of these CNVs clonally (**Fig. 3h**). Among these, gains of Chromosome 17 or 17q were the most frequent aberrations, followed by gain of whole Chromosome 7 and LOH of whole Chromosome 8. In cases with detectable ECA, these CNVs were frequently associated with the ECA, suggesting that they might be the earliest events during Group 3/4 medulloblastoma initiation. In contrast to the high abundance of CNVs, focal events in known driver genes (SNVs, indels, focal amplifications/deletions or structural rearrangement) were overall rare (**Fig. 3h**). Among these, amplification of *MYC* or *MYCN* and duplication of *SNCAIP* leading to *PRDM6* overexpression were the most frequent alterations (**Fig. 3h**). However, the single-cell analysis (c.f. **Fig. 2b,f,j**), reveals that these mutations were mostly subclonal. Interestingly, Group 3/4 medulloblastomas with amplification of *MYC* or *MYCN*, or duplicated *SNCAIP* had significantly higher SNV densities at both ECA (**Fig. 3i**) and MRCA (**Fig. 3j**) than the remaining tumors, suggesting that later onset of (pre-)malignancy may predispose to the subsequent acquisition of these drivers. When quantifying the burden of focal driver mutations overall, Group 3/4 medulloblastomas without a detectable ECA had on average no focal driver mutation at clonal variant allele frequency (VAF) as compared to two large-scale clonal CNVs (**Fig. 3k**). Group 3/4 medulloblastomas with a detectable ECA had significantly more clonal driver CNVs than cases without (p = 0.04, Wilcoxon Rank Sum Test); these cases harbored with an average of four clonal driver CNVs as opposed to no focal driver mutation at clonal VAF (**Fig. 3l**). In contrast to the stark difference between CNVs and focal mutations among clonal drivers, the majority of cases with (62%) and without (73%) detectable ECA had no subclonal driver at all with no significant difference between CNVs and focal driver mutations (**Fig. 3k,l**). Thus, the mutational landscape in Group 3/4 medulloblastoma suggests a fundamental role of CNVs during tumor initiation, while additional mutations acquired during disease progression seem to drive additional evolution in a subset of tumors only.

### Subclonal spatial heterogeneity: interspersed or segregated

To better understand the spatial relationship of the tumor subclones and reveal further insight into their evolution, we performed spatial transcriptomics on samples with available material (n=13 primary, n=4 relapse). We used technology that applies multiplexed *in situ* hybridization of a selected gene set to achieve single-cell spatial resolution of a tumor sample (**Extended Data Tables 4**,**5**). UMAP visualization of merged spatial data from primary tumor samples reflected combined snMultiomic profiling, allowing us to distinguish subgroup-specific properties and identify non-tumor cell types (**Extended Data Fig. 5a-e**). The spatial locations of tumor cells were determined by the expression of *MYC, MYCN*, and *PRDM6* (**Fig. 4a**) along with other genes associated with proliferation (*e.g. MKI67*, **Extended Data Fig. 5f**). The tumor microenvironment, including glial, immune and meningeal cells, was characterized using cell type-specific markers (**Extended Data Fig. 5f**).

**Figure 4.**
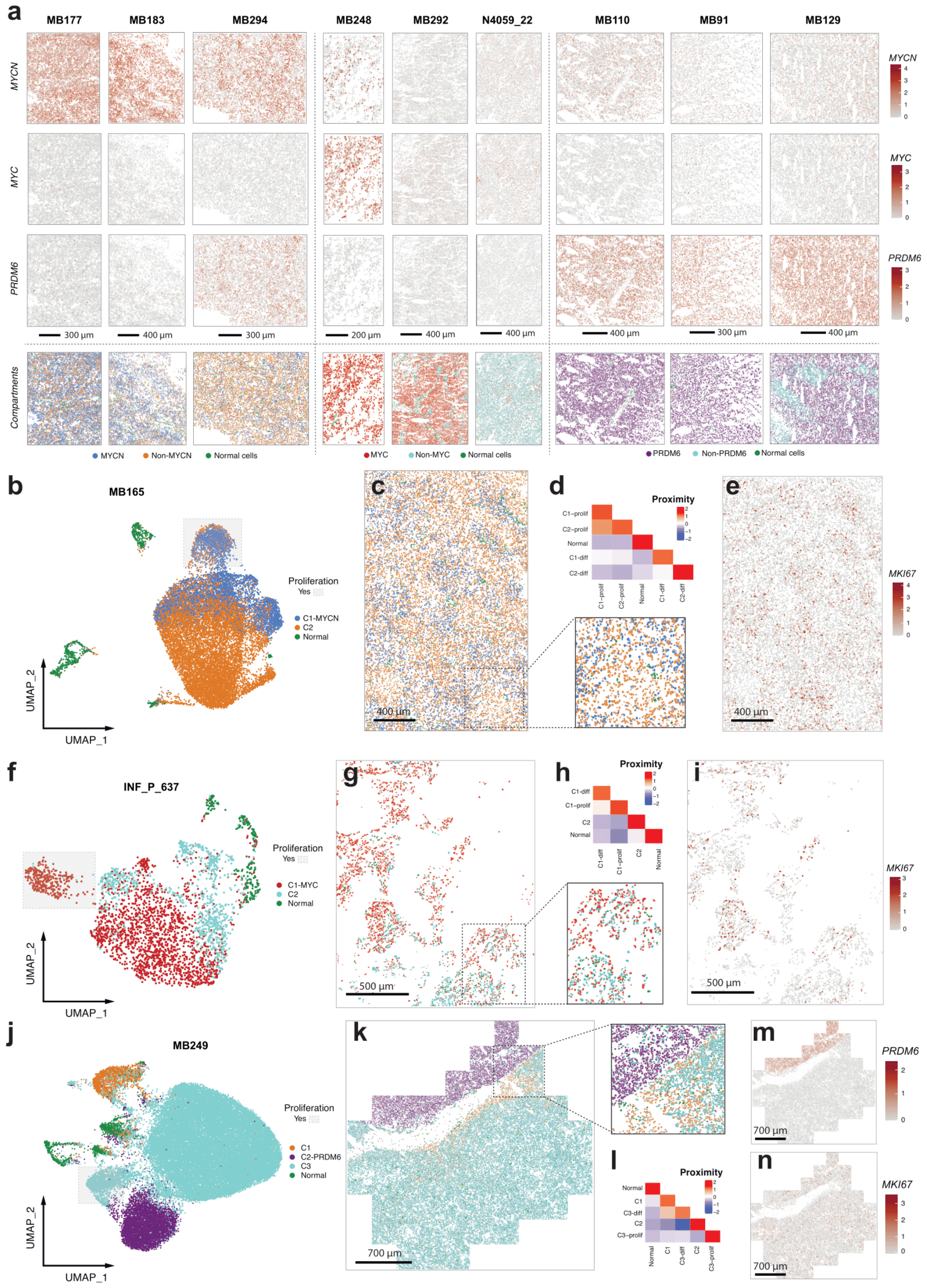
Spatial heterogeneity across oncogene-present Group 3/4 medulloblastoma samples. a) Spatial gene expression of *MYC, MYCN, PRDM6*. Last row, projection of clones derived from snRNA-seq. b) Spatial data UMAP of representative *MYCN* sample. c) Spatial visualization of clones of sample in b. Enlarged view of a fragment in the bottom right. d) Proximity of each compartment to each other of sample in b. e) *MKI67* spatial expression of sample in b. f) Spatial data UMAP of representative *MYC* sample. g) Spatial visualization of clones of sample in f. Zoom into specific region on bottom right. h) Proximity of each compartment to each other of sample in f. i) *MKI67* spatial expression of sample in f. j) Spatial data UMAP of representative *PRDM6* sample. k) Spatial visualization of clones of sample in j. Zoom into specific region on top right. l) Proximity of each compartment to each other of sample in j. m) *PRDM6* and n) *MKI67* spatial expression of sample in j.

To determine the spatial distribution of the identified subclones, we projected the snRNA/ATAC data onto the spatial data (**Fig. 4a**, last row). Overall, we distinguished two major spatial localization patterns: *interspersed*, where independent subclones mixed throughout the tumor sample, and *segregated*, where a clear boundary between independent subclones could be delineated. In the majority of cases, the subclones were interspersed, as observed from the spatial distribution of the corresponding marker gene expression.

In *MYCN-*amplified tumors, the observed clonal architecture derived from snRNA-seq data was also present in the spatial data (**Fig. 4b**). The subclones exhibited an interspersed spatial pattern (**Fig. 4c**), with pockets of MYCN and non-MYCN clones highlighted through neighborhood enrichment within the tumor sample (**Fig. 4c**, inset). The proliferating compartments within these subclones were also interspersed across the tumor tissue observed by *MKI67* expression (**Fig. 4e, Extended Data Fig. 5f**). On a smaller scale, however, neighborhood enrichment analysis showed that proliferating cells of subclones clustered together, away from the differentiating cells **(Fig. 4d**). The normal cells were mostly isolated from the tumor subcompartments.

A similarly interspersed spatial pattern of MYC and non-MYC subclones was present in *MYC*-amplified tumors (**Fig. 4f,g**), yet islands of segregated non-MYC subclones were also observed (**Fig. 4g**, inset). Proliferating cells (*MKI67*+) were interspersed throughout the tumor tissue (**Fig. 4i, Extended Data Fig. 5f**). The differentiated tumor cell compartment within the *MYC*-subclone (C1-diff) was in closer proximity to the proliferating cell compartment within the same subclone (C1-prolif, **Fig. 4h**). Normal cells were largely isolated from the *MYC-*amplified subclone compartments.

We observed a segregated spatial separation of subclones in the *PRDM6* sample (**Fig. 4j,k,m**). This spatial segregation of clones was confirmed in another region of the same tumor specimen (**Extended Data Fig. 5g,h**). While the subclones were segregated, the proliferating cell compartments within the subclones were interspersed within the spatial block (**Fig. 4n, Extended Data Fig. 5i**). As expected, the neighborhood enrichment analysis in this sample showed the separation of *PRDM6* clone from other compartments (**Fig. 4I**). In another sample we also confirmed the dual-oncogene, PRDM6-MYCN subclone (**Fig. 2b**, bottom right) where cells with activity in both genes reside in the same spatial regions (**Extended Data Fig 5k-l**).

Altogether, the observed spatial patterns suggest that clonal evolution does not lead to spatial compartmentalization within tumors. Instead, cellular migration may dictate communication among the clones, which can then drive competing or collaborative interactions among the co-existing tumor populations.

### *MYC*-driven subclones take over during tumor evolution

We identified a primary tumor sample with two distinct subclones (MB272), harboring *MYC* (C3) and *MYCN* (C2) amplifications simultaneously (**Fig. 2f, 5a-c**). This subgroup II sample was originally characterized as *MYC-*amplified only, while *MYCN* amplification was not clearly observed in the bulk DNA methylation CNV profile (**Extended Data Fig. 6a**). The *MYC* and *MYCN*-subclones in this tumor sample had proliferating and differentiating compartments (**Fig. 5b, c**). The *MYC*-subclone was strongly enriched with progenitor-like activity (**Extended Data Fig. 6b**). Remarkably, a clear spatial separation was observed between *MYC-* and *MYCN-* expressing cells (**Fig. 5d; Extended Data Fig. 6c**,**d**). This spatial segregation (**Fig. 5e,f**) reflected the phylogenetic tree of tumor evolution projected from the snRNA-seq CNV annotation (**Fig. 2f**). Additional sets of subclone-specific genes also showed explicit spatial specificity (**Extended Data Fig. 6e,f**). As expected, low contact proximity was identified between *MYC* and *MYCN* subclones (**Extended Data Fig. 6g**).

**Figure 5.**
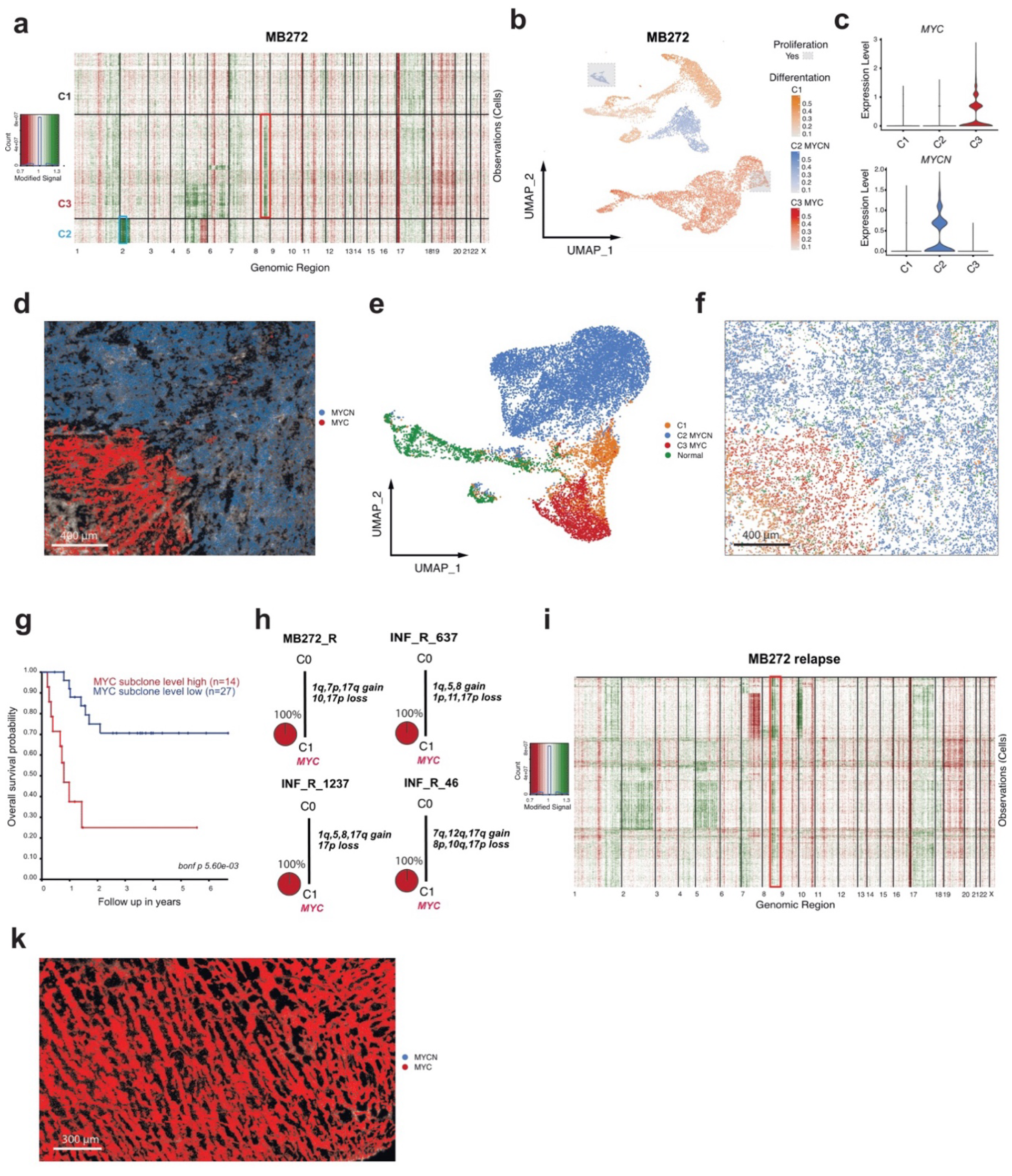
Independent oncogene subclones may co-occur in one tumor, but subclones are lost at relapse. a)Copy number profiles of snATAC-seq data from *MYC*-*MYCN* sample MB272. Red, chromosome loss. Green, chromosome gain. b) snRNA-seq UMAP of sample shown in b. Gray boxes, proliferating cells. Blue, differentiation signal enrichment in *MYCN*-expressing C2 clone. Red, differentiation signal enrichment in *MYC*-expressing C3 clone. Orange, differentiation signal enrichment in C1 clone. c) *MYC* and *MYCN* expression in C1, C2, and C3 clones. d) Spatial gene expression of *MYC* (red) and *MYCN* (blue) from original signals. e) Spatial data UMAP of sample shown in d. f) Spatial visualization of clones of sample in d. g) Kaplan–Meyer overall survival probability curves for medulloblastoma Subgroup V tumors with high (red) and low (blue) *MYC* subclone level enrichment. h) Somatic phylogeny trees for *MYC* relapse samples. i) Copy number profiles of snATAC-seq data of relapse sample arising from primary sample shown in a-f. k) Spatial gene expression of *MYC* (red) and *MYCN* (blue) in spatial transcriptomic relapse sample of case shown in a-f.

According to current knowledge, *MYC* and *MYCN* amplifications are considered mutually exclusive events^25^ in medulloblastoma (**Extended Data Fig. 6h**) and other tumor types^26^. Because of the unexpected occurrence of both amplifications in this case, we conducted a systematic analysis across a larger medulloblastoma cohort to identify additional cases where these oncogenes may co-occur. We identified 6 putative cases based on DNA methylation CNV profiles (**Extended Data Fig. 6i**). Using immunohistochemistry, we identified another case where both MYC and MYCN staining were seen in the same tumor sample (**Extended Data Fig 6j**). During the preparation of this manuscript, a case study reported a primary tumor sample where both *MYC* and *MYCN* amplified cells were present^27^. Altogether these independent cases suggest the possibility that *MYC* and *MYCN* amplifications co-occur within the same tumor more frequently than originally thought. Nevertheless, it is noteworthy that individual cells within the tumor express only one of these oncogenes and are spatially segregated.

The presence of *MYC* and *MYCN* subclones cannot be distinguished using the bulk profile techniques due to potential low presence of cells of a particular subclone in the obtained data. Therefore, we generated unique signatures of *MYC-* and *MYCN-* subclones derived from single-cell data to identify additional samples harboring two oncogene amplifications. We performed a deconvolution analysis of bulk transcriptome profiles, using the *MYC*/*MYCN* case as the reference control. Using this method, we detected additional samples where *MYC* and *MYCN* subclones may co-occur in the same sample (**Extended Data Fig. 6k**). We validated this finding using FISH on an identified sample with available material (**Extended. Data Fig. 6l**).

We next checked whether this information could be exploited for diagnostic purposes. Therefore, we investigated whether the relative presence of *MYC* or *MYCN* subclones derived from the deconvolution analysis predicted patient outcomes. In subgroup V, 4 out of 41 cases harbored a known *MYC* amplification, based on CNV profiles, and correlated with a low probability of survival (**Extended Data Fig. 6m**). We identified 4 potential cases with an occurrence of a *MYC*-amplified subclone based on deconvolution. These patients had a lower overall survival (**Fig. 5g**). Therefore, the poor outcomes of subgroup V patients may be explained by an undiagnosed *MYC* subclone that potentially outcompetes other subclones to drive relapse.

To further test this possibility, we performed single-nucleus molecular profiling on 4 relapse *MYC-*amplified cases. In all relapse cases, new subclones arose, but all tumor cells harbored the *MYC* amplification (**Fig. 5h,i**). For example, in the matched relapse *MYC/MYCN* case, the *MYCN* subclone is lost at relapse (**Fig. 5i**). This loss of *MYCN* expression was confirmed using spatial transcriptomics (**Fig. 5k, Extended Data Fig. 6n,o**). These results suggest that the *MYC* subclone outcompetes other subclone(s) during tumor progression and hence the presence of subclonal *MYC* amplification at diagnosis may predict the probability of relapse.

## Discussion

Despite advances in understanding the cellular origin of Group 3/4 medulloblastoma, the tumor initiating and driving mechanisms remain elusive. We show that the genetic aberrations that lead to overexpression of oncogenes are not the initiating events in Group 3/4 medulloblastoma. Therefore, *MYC* and *MYCN* are likely not the primary “drivers”, but instead are acquired after malignant transformation and likely accelerate tumor growth. Instead, our results suggest that the initiating or ‘driving’ events in Group 3/4 medulloblastoma are large-scale CNVs, likely occurring during fetal development. This finding is in line with the hypothesis that tetraploidization is a frequent early event in medulloblastoma^10^ and is associated with survival rates and high risk of relapse^28^. Intriguingly, another pediatric tumor, neuroblastoma, has a similar order of genetic events: CNVs are the initiating event and occur early in the first trimester of pregnancy^18^. How exactly large-scale CNVs drive early tumorigenesis in different cell types and whether this knowledge can potentially be exploited for early cancer detection remains to be explored.

Group 3/4 medulloblastoma with *MYC, MYCN*, or *PRDM6* alterations have complex subclonal structure, with each subclone having unique properties. Therefore, our results strongly argue against the dogmatic “cancer stem cell” hierarchy for Group 3/4 medulloblastoma, as these tumors maintain distinct subclones with separate proliferating and differentiating compartments, irrespective of the presence of recognized oncogenes.

In addition, undetected *MYC*/*MYCN* subclones challenge the bulk analysis approach in the diagnostic space. These findings, along with support from others^29^ suggest that single-cell analyses may be an important diagnostic tool in the future, especially for Group 4 and subgroup V tumors. In addition, our data challenge the cutoffs used for a tumor to be called *MYC*/*MYCN*-amplified by FISH, as even the smallest *MYC* subclone, which initially are lowly abundant, have the potential to expand into the dominant clone during relapse. *MYC* amplification may drive disease progression and contribute to therapy resistance and relapse. Such a pattern of *MYC* dominance in subclonal evolution was observed in other tumors including gliomas^30^, suggesting that our results may be relevant also to other tumor entities associated with *MYC* oncogenesis.

## Extended Data Figures

**Extended Data Figure 1.**
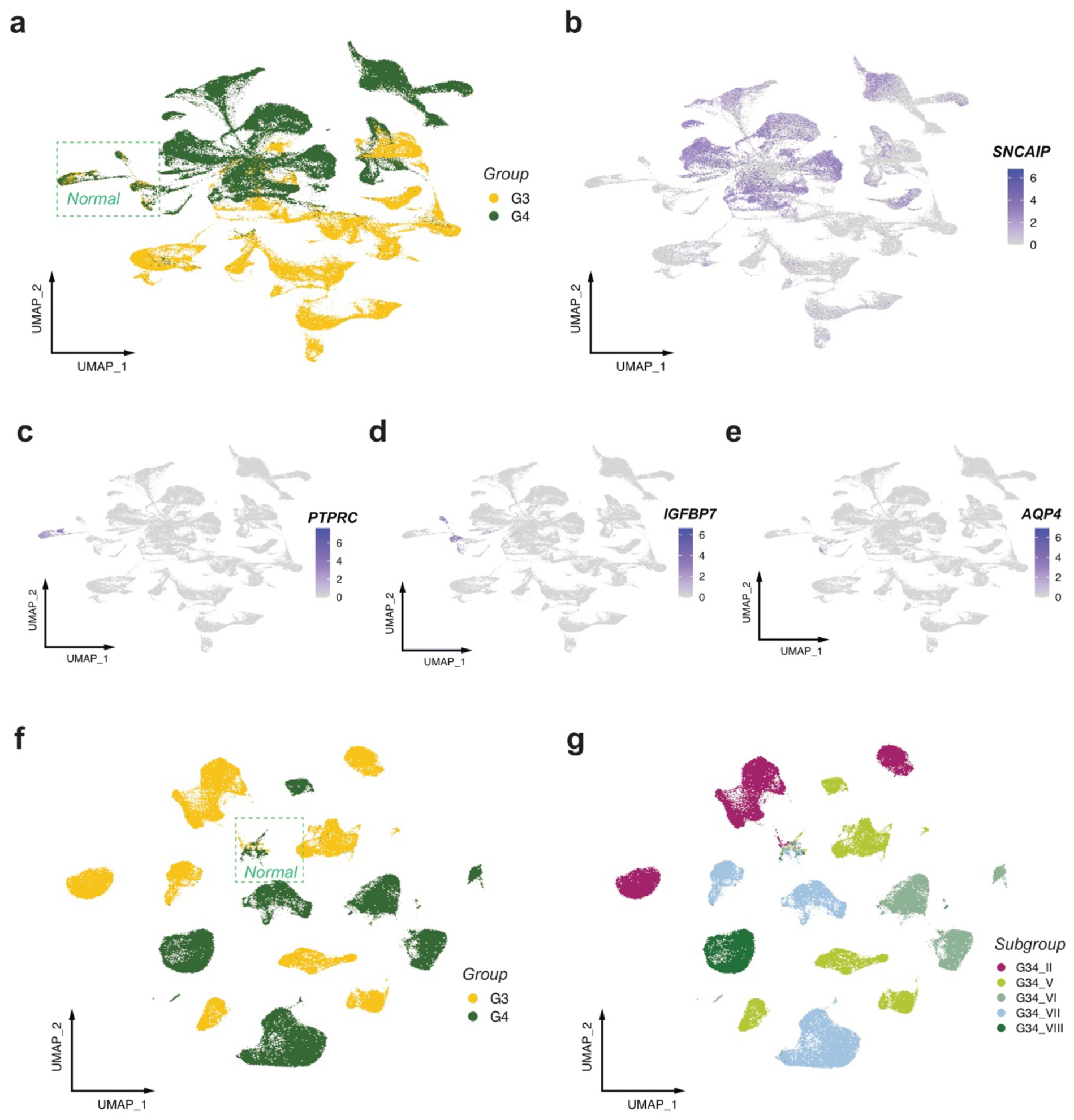
Group 3/4 Medulloblastoma single nuclei RNA and ATAC profile properties. a) UMAP of snRNA-seq merged dataset with MB groups annotation. Non-tumor cells marked. Feature plots showing a)*SNCAIP*, c) *PTPRC*, d) *IGFBP7* and e) *AQP4* and expression within UMAP of merged snRNA-seq dataset. f) UMAP of snATAC-seq merged dataset with medulloblastoma groups annotated. Green box, normal cells. f) UMAP of snATAC-seq merged dataset with medulloblastoma subgroups annotated.

**Extended Data Figure 2.**
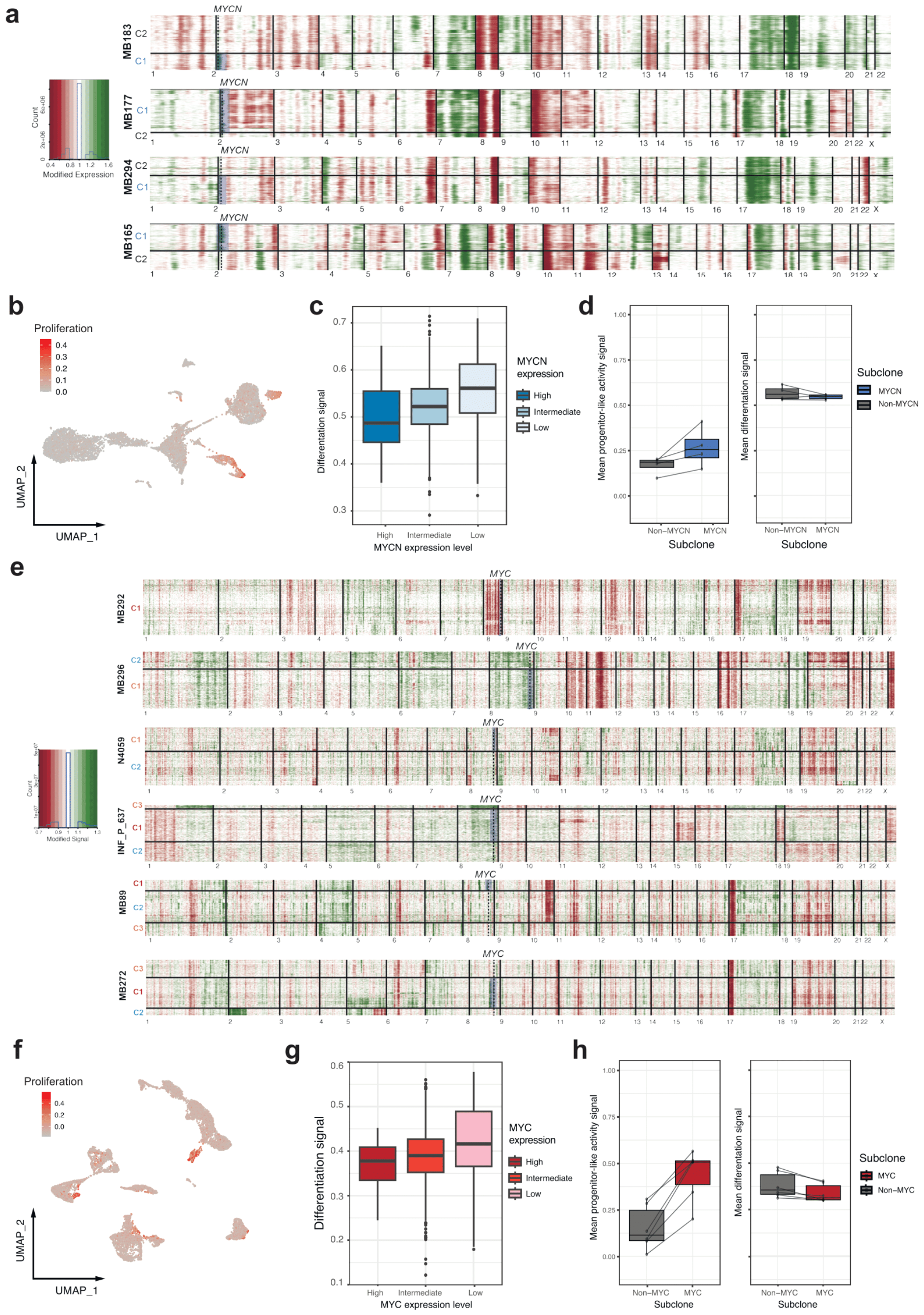
Copy number profiling of single nuclei profiles from Group 3/4 MYC-and MYCN-amplified samples. a) Copy number profiles of snRNA-seq data from *MYCN* samples (n=4). b) Per cell GSVA enrichment of proliferation markers within UMAP of MB183 *MYCN-*amplified sample. c) Differentiation signal compared to ranked *MYCN* expression within *MYCN*-amplified subclone in sample MB183. *MYCN* normalized expression cutoffs: low = zero, intermediate > 0 and < 2, high > 2. d) Boxplots showing difference in mean signal of progenitor-like activity (left side) and differentiation (right side) between *MYCN*-amplified and non-*MYCN*-amplified subclones in n=4 tumor cases. e) Copy number profiles of snATAC-seq data from *MYC* samples (n=6). f) Per cell GSVA enrichment of proliferation markers within UMAP of MB89 *MYC-*amplified sample. g) Differentiation signal compared to ranked MYC expression within MYC-specific sublclone in sample MB89. *MYC* normalized expression cutoffs: low = zero, intermediate > 0 and < 2, high > 2. h) Boxplots showing difference in mean signal of progenitor-like activity (left side, t-test p-val: 0.003) and differentiation (right side) between *MYC*-amplified and non-*MYC*-amplified subclones in n=6 tumor cases.

**Extended Data Figure 3.**
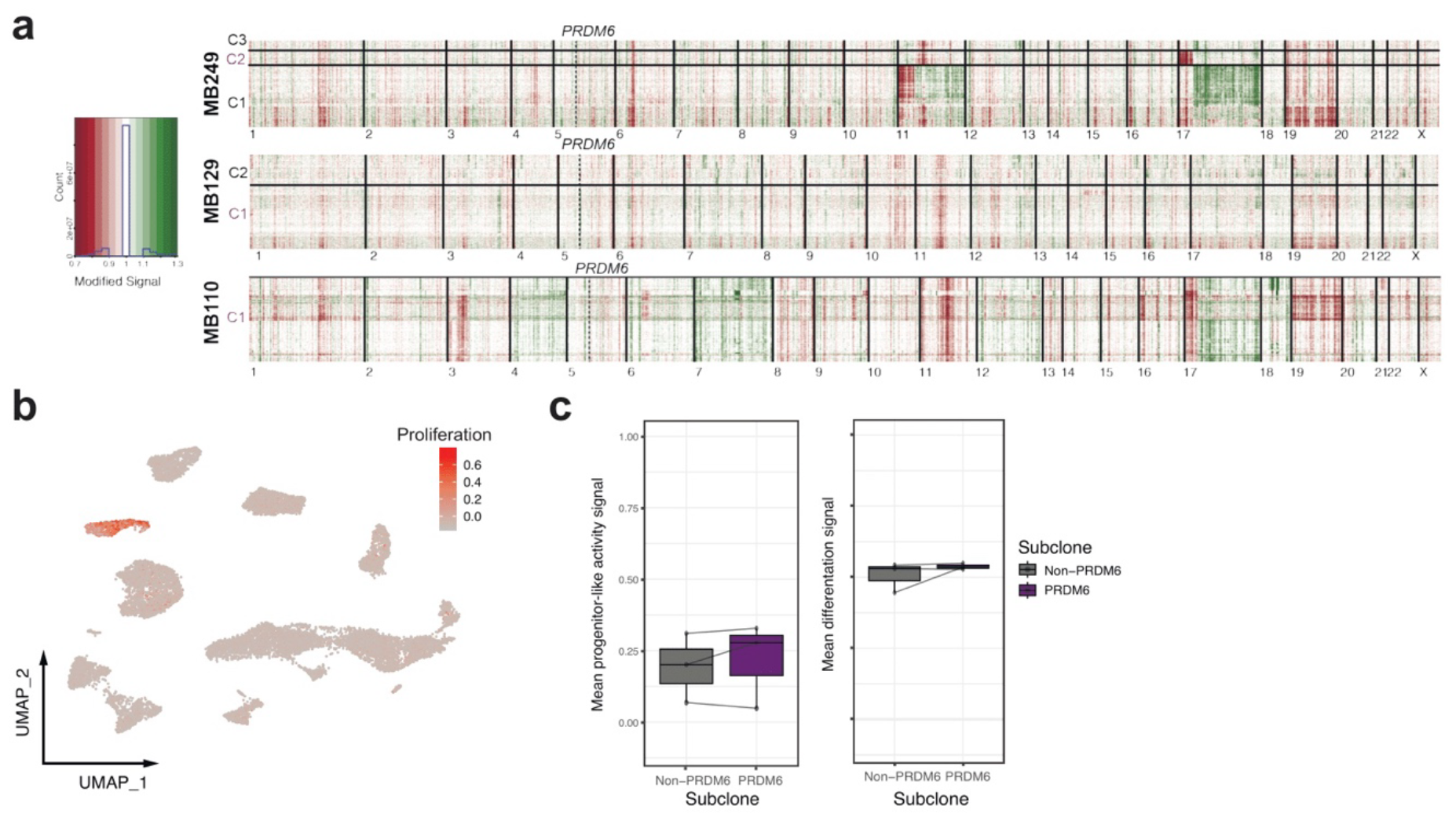
Copy number profiling of single nuclei profiles from Group 3/4 PRDM6 samples. a) Copy number profiles of snATAC-seq data from *PRDM6* samples (n=3). b) Per cell GSVA enrichment of proliferation markers within UMAP of MB249 *PRDM6* sample. c) Boxplots showing difference in mean signal of progenitor-like activity (left side) and differentiation (right side) between *PRDM6*-and non-*PRDM6* subclones in n=3 tumor cases.

**Extended Data Figure 4.**
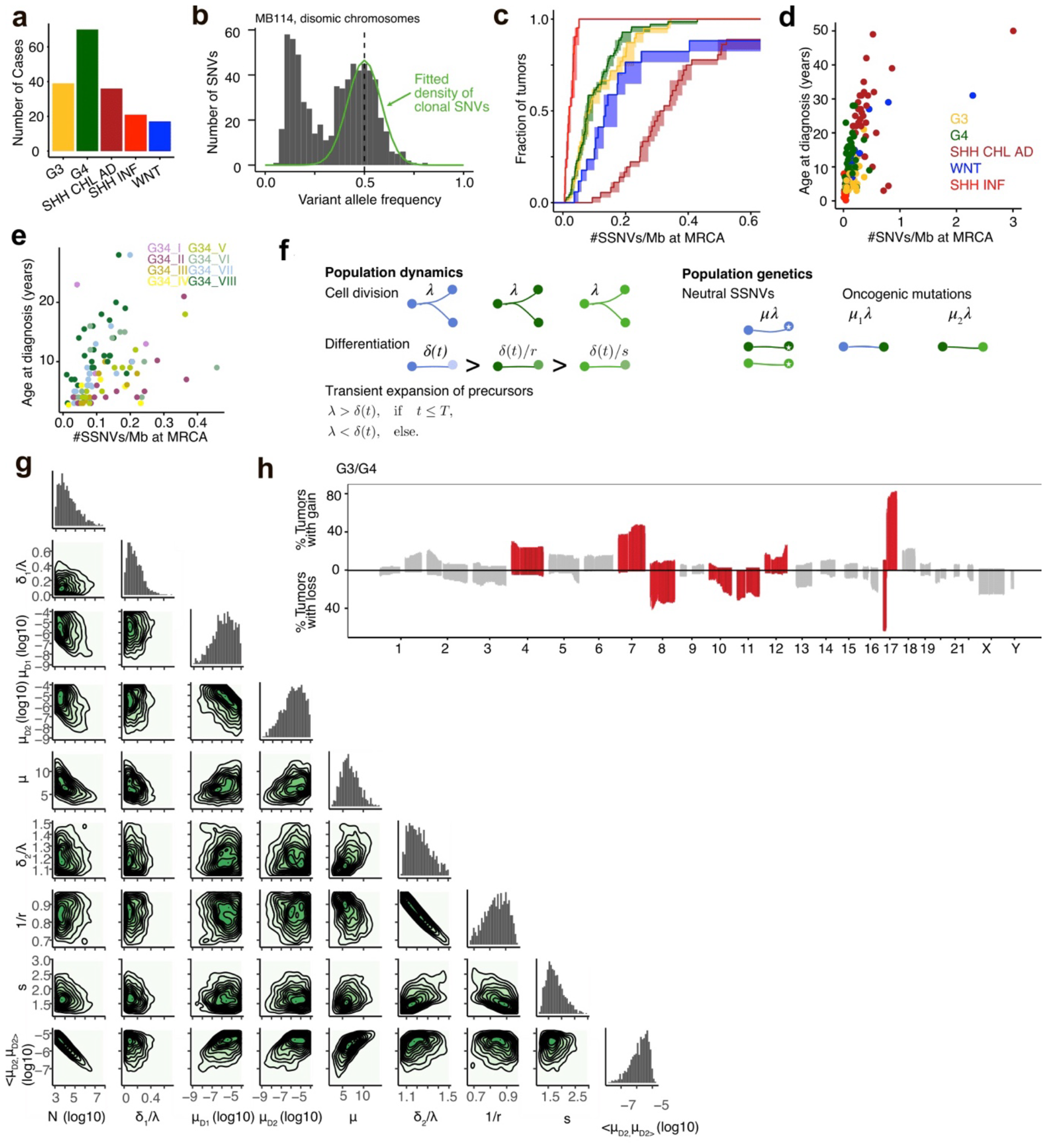
Group 3/4 medulloblastoma somatic mutation profiles. a) Overview of medulloblastoma samples analyzed by bulk WGS and stratified by subgroup. b) Variant allele frequencies of SNVs detected in sample MB104 (G34_VIII) on disomic chromosomes. Green line, fitted density of clonal SNVs; dashed vertical line, true clonal VAF according to the sample purity (as estimated by ACEseq). c), SNV densities at MRCA stratified by subgroup (densities were quantified from 39 MB G3, 70 MB G4, 21 MB SHH INF, 36MB SHH CHL/AD, and 17 MB WNT; 4 MB SHH CHL/AD and 4 MB WNT stood out with clonal mutation densities between 0.7 and 2.9 SNVs/Mb and are not shown). Mean and 95% CI, estimated by bootstrapping the genomic segments 1,000 times. d) Mean SNV densities at MRCA versus age at diagnosis across groups (n = 175 cases with age information). e) Mean SNV densities at MRCA vs age at diagnosis across G3/4 subgroups (n = 106 cases with age information). f) Population genetics model of tumors initiation in two steps of clonal selection. Cells in the tissue of origin divide at rate *λ* and differentiate at rate *δ*. The tissue initially expands until time *T* and thereafter contracts. Somatic variants are acquired at rate *μλ* and driver mutations are acquired at rate *μ*_*1*_*λ* and *μ*_*2*_*λ*, respectively. The latter decrease the differentiation rate by a factor *1/r* and *1/s*, respectively. g) One-dimensional and two-dimensional posterior probabilities for the model fit to all G3/4 MBs (n = 109). < *μ*_*1*,_ *μ*_*1*_*>* denotes the geometric mean of the driver mutation rate. h) Percentage of tumors with copy number gains and losses ≥1Mb along the genome. Red, Regions where CNVs were significantly more frequent than expected, according to a Binomial test with p_adj_ < 0.01; Holm correction for multiple sampling.

**Extended Data Figure 5.**
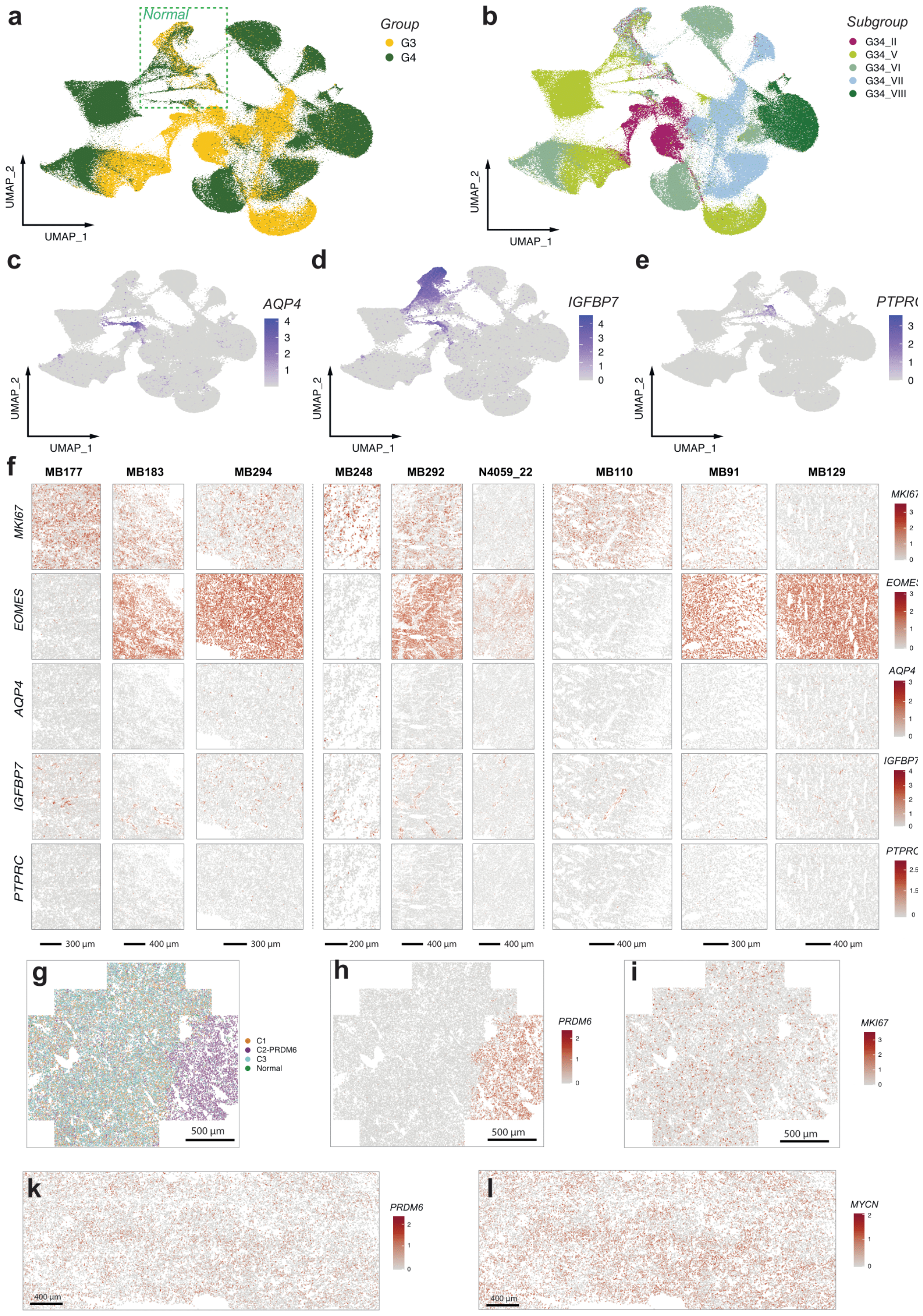
Spatial resolution of sub-clonal tumor populations. a) UMAP of spatial merged dataset with MB groups annotation. Normal cells marked. b) UMAP of spatial merged dataset with MB subgroups annotation. c-e) Feature plots showing c) *AQP4*, d) *IGFBP7* and e) *PTPRC* expression within UMAP of merged spatial dataset. f) Spatial gene expression of *MKI67, EOMES, AQP4, IGFBP7* and *PTPRC* across samples. g) Spatial visualization of clones of PRDM6 sample in 2^nd^ image fragment. h) *PRDM6* and i) *MKI67* spatial expression of sample in g. k) *PRDM6* and l) *MYCN* spatial gene expression in image fragment of sample MB292.

**Extended Data Figure 6.**
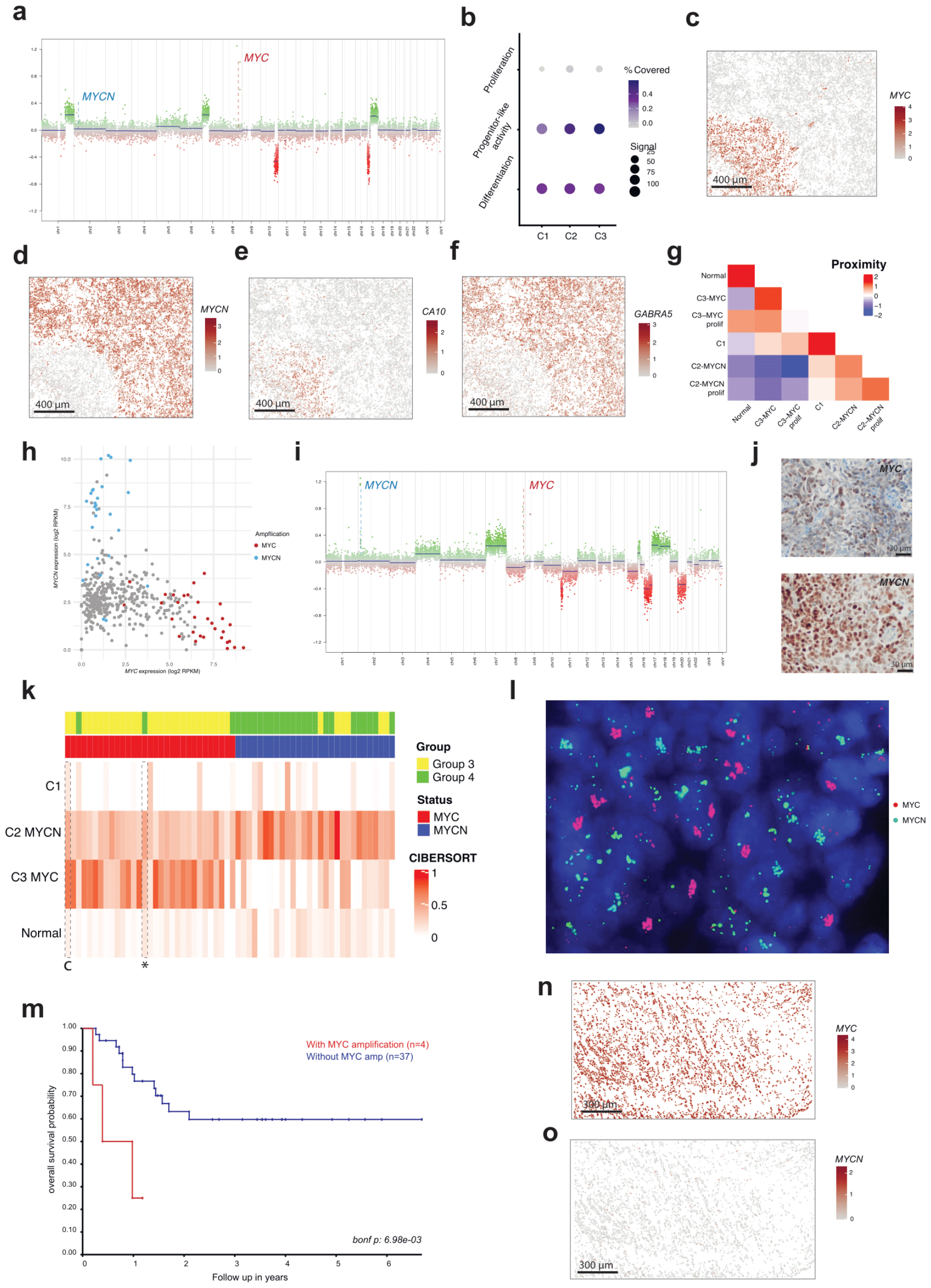
Independent oncogene clones may co-occur in one tumor. a) CNV profile of MB272 cases bulk methylation data. b) Per cell GSVA enrichments of proliferating, progenitor-like and differentiation in single sample MB272. Spatial expression of c) *MYC*, d) *MYCN*, e) *CA10* and f) *GABRA5* in sample MB272. g) Proximity of each compartment to each other in sample MB272 spatial data. h) Negative correlation (R=-0.287, p=1.08e-09) between *MYC* and *MYCN* expression within medulloblastoma FFPE bulk RNA-seq cohort (n=435). i) CNV profile of bulk methylation data from a Group 3/4 tumor with amplifications of *MYC* and *MYCN*. j) Identification of MYC (left) and MYCN (right) signals in the same sample using IHC. k) CIBERSORT deconvolution results across subset of *MYC*/*MYCN* cases from medulloblastoma bulk FFPE RNA-seq cohort. MB272 single cell data with subclones annotation used as a reference, the data from control case is marked with c, target sample marked with asterisk. l) Identification of *MYC* (red) and *MYCN* (green) signals in the highlighted target Group 3/4 sample described in panel (k) using FISH. m) Kaplan–Meyer overall survival probability curves for medulloblastoma Subgroup V tumors with (red) and without (blue) *MYC* amplification as identified from bulk data CNV profiling. n) *MYC* and o) *MYCN* expression in spatial transcriptomic relapse sample.

## Extended Data Tables

**Extended Data Table 1**. Overview of target medulloblastoma Group 3/4 cohort with focus on snRNA-seq, snATAC-seq and single cell spatial transcriptomics data.

**Extended Data Table 2**. Differentially expressed genes, specific for *MYC, MYCN* and *PRDM6* subclones confirmed in minimum n=3 samples.

**Extended Data Table 3**. Overview of target medulloblastoma ICGC cohort with focus on WGS data.

**Extended Data Table 4**. List of 100 target genes applied for the spatial single cell protocol.

**Extended Data Table 5**. Quality control overview of spatial single cell data.

## Methods

### Target cohort selection and verification

Target tumor tissue samples were collected from medulloblastoma global published materials (ICGC^3^, FFPE^11^ and INFORM^31^ cohorts) and are described further in Joshi *et al bioRxiv*. For each selected case the copy number/structural variant profiles from methylation and/or whole genome sequencing data were used to identify *MYC*/*MYCN* amplification and *SNCAIP* structural variant presence. Bulk gene expression RNA-seq profiles from these samples were used to inspect *MYC/MYCN/SNCAIP/PRDM6* expression as well. For some cases with sufficient available FFPE material, additional FISH experiments were performed to verify the selection (details in **Extended Data Table 1**). No statistical methods were used to predetermine sample size.

### Single-nuclei multi-omics sequencing

Flash frozen tumor samples were processed to extract nuclei as described earlier^23^, and described in detail in Joshi *et al bioRxiv*. Extracted nuclei were processed using Chromium Single Cell Multiome ATAC Gene expression kit and Chromium Controller instrument (10x genomics) as per manufacturer’s recommendations. One sample, MB248, was processed with Chromium Next GEM Single Cell 3’ reagent kit as per manufacturer’s recommendation. 15,000-20,000 nuclei were loaded per channel along with the multiome gel bead. Libraries were quantified using Qubit Flurometer (Thermo Fisher Scientific) and profiled using Fragment Analyzer. GEX and ATAC libraries were sequenced using NextSeq2000 to recommended lengths. If ATAC library was not of good quality, we still used obtained RNA library if that was found to be appropriate based on QC parameters. RNA-seq and ATAC-seq datasets were further analyzed separately.

### Single nuclei RNA sequencing data analysis

De-multiplexed reads were aligned to human genome assembly GRCh38 (v. p13, release 37, gencodegenes.org). Comprehensive gene annotation (PRI) was customized by filtering to transcripts with following biotype: protein coding, lncRNA, IG and TR gene and pseudogene as recommended for *cellranger mkgtf* wrapper. Reads were aligned using STARsolo with parameters: --soloType *CB_UMI_Simple* --soloFeatures *Gene GeneFull* --soloUMIfiltering *MultiGeneUMI* -- soloCBmatchWLtype *1MM_multi_pseudocounts* --soloCellFilter *None* --outSAMmultNmax *1* -- limitSjdbInsertNsj *1500000*. For overlapping genes where intronic alignment recovered low counts, exonic alignment counts were used. Cells were separated from debris using *diem* pipeline^32^. Cells with mitochondria fraction above 2%, number of detected genes above 6600 and an intronic fraction (number of reads aligned to intron/total number of reads aligned to exon+intron) less than 25% were also filtered out. Filtered cells were corrected for background signature using SoupX pipeline^33^. Finally scrublet^34^ tool was used to remove putative doublets. Further the gene expression matrices from all samples were merged together in full matrix and processed via Seurat package^35^ to normalize, compute top principal complements (n=30), find most highly variable genes (n=2500) and visualize via UMAP. After distinguishing non-tumor cells based on corresponding markers and combined UMAPs, per sample processing was performed using Seurat toolkit using same settings combined with cells clustering. The enrichment of proliferation, differentiation and progenitor-like activity of medulloblastoma-specific markers per cell was performed using single sample function from GSVA R package^36^ using two independent reference datasets^6,7^.

### Single nuclei ATAC sequencing data analysis

ATAC reads were aligned to GRCh38 using Cellranger arc wrapper. The selected cells were processed using Signac R package^37^ to filter the doublets/outliers based on signal per cell distribution analysis and inspect the cell compartments via UMAP visualization after normalization and identification of most highly-variable regions. Single nuclei derived RNA-seq data information was used to annotate the cells from corresponding processed data.

### Single cell CNV phylogeny reconstruction

Before CNV analysis transferring of single cells into meta-cells was performed by computing sum of gene expression counts across n=5 cells combined within the clusters derived from initial Seurat processing in order to improve the specificity. For snRNA-seq data all filtered genes were used as input matrix for InferCNV tool^12^ using droplet protocol adjusted parameters and hierarchical clustering to derive potential phylogeny. Normal cells identified previously were used as reference control for each sample. After CNV calling further the initial clustering, *MYC/MYCN/PRDM6* expression as well as computed per cell progenitor-like activity enrichment values were used to finalize the derived phylogeny for each case from manual inspection. Differentially expressed genes for identified subclones per sample were computed via Wilcoxon Rank Sum test.

For snATAC-seq data a matrix with all genomic regions as raw and read counts per column per sample were used to adjust for InferCNV input format. Further meta-cells formation and same CNV calling procedure as for snRNA-seq were performed on the derived matrices. The subclone annotation derived from snRNA-seq data was used to assign corresponding clusters phylogeny per sample.

### Molecular Cartography (MC)

Specific gene set (n=100) covering Group 3/4 known driver genes alongside marker genes of the developing cerebellum non-malignant cell types was selected for the protocol (**Supplementary Table 4**). OCT-embedded samples were cryo-sectioned into 10 μm sections onto an MC slide. Fixation, permeabilization, hybridization and automated fluorescence microscopy imaging were performed according to the manufacturer’s protocol (Molecular preparation of human brain (beta), Molecular coloring, workflow setup) as described previously^38^.

### Spatial data analysis

The detection of cell boundaries was performed with CellPose^39^. Afterwards, gene expression counts were computed per cell and extracted using additional custom Python scripts. Initial cells filtering was performed by assigning minimum number of counts/genes per cell and size of the cells. Afterwards, the analysis of the formed gene expression matrix, including clustering and UMAP visualization, was executed using the Seurat toolkit^35^. Annotation of cell states and types was achieved through direct projection with the snRNA-seq data via transfer function and verified from visual inspection of marker genes. Spatial-specific analysis, the detection of closest cell connections, was conducted using the Giotto toolkit^40^.

### Deconvolution analysis of MYC/MYCN cases

The assigned *MYC/MYCN* single cell dataset (MB272) with annotation of compartments was used as reference control for the CIBERSORT method^41^ in order to perform deconvolution on a set of bulk FFPE medulloblastoma RNA-seq profiles from *MYC/MYCN* samples^11^. For each case, *MYC/MYCN* status was derived from methylation copy number profile. The obtained deconvolution values were visualized via ComplexHeatmap R package to correspond compartment enrichments per sample to *MYC* and *MYCN* status.

Survival analyses based on the expression of *MYC* as well as computed deconvolution *MYC* compartment proportion multiple genes were performed using Kaplan-Meyer algorithm with applied Bonferroni correction for multiple testing. The result plots were generated via R2 Genomics Analysis and Visualization Platform (http://r2.amc.nl).

### Whole genome sequencing data

Mutation calls (SNVs, indels, CNVs and SVs) of previously published whole genome sequencing data from medulloblastoma of all subtypes were taken from ICGC dataset^3^. Only samples from primary tumors with clear subtype annotation and clear ploidy status were included; see **Extended Data Table 3** for an overview on these samples and associated clinical data.

### Driver mutations (SNVs and CNVs)

Nonsynonymous SNVs, small indels and small structural rearrangements (amplifications, defined as copy number gains ≥10, homozygous deletions with <0.9 copy numbers, and translocations with a minimal event score of 5) were classified as driver mutations if they targeted a splice-site or an exonic region of *PRDM6, MYC*, or a gene listed as a putative driver of medulloblastoma in the cancer driver database intogen^42^ (release date 23/05/31). Moreover, we included *TERT* promoter mutations at hg19 positions 1295228 and 1295250 as drivers. High-level amplifications affecting *MYC* or *MYCN* (identified from methylation/WGS copy number profiles) and duplications of *SNCAIP*, leading to overexpression of *PRDM6* (identified from WGS SV calling) were additionally integrated from previous global data analysis^3^.

Large-scale copy number variants (CNVs) were defined as CNVs spanning at least 1Mb and with a coverage ratio < 0.9 or a coverage ratio > 1.1, according to the output by ACEseq. Retained CNVs with a size of at least 25% the size of the p arm of a respective chromosome were further classified as affecting both arms if the CNV spanned the centromere, and else as affecting the p arm or q arm. Among these CNVs, we tested for positive enrichment of particular chromosomes in the cohort using a binomial test with success probability 1/24 (i.e., assuming that each chromosome has equal probability to be affected by the CNV). Chromosomes with an adjusted p value < 0.05 (Holm’s correction) were classified as likely drivers of medulloblastoma. This analysis was separately performed for gained and lost chromosomes. Among G3/4 medulloblastomas, we identified gains of Chromosome 4, 7/7q, 12, 17/17q and losses of Chromosome 8, 10/10q, 11 as significant. We augmented this list by gains of Chromosome 18, 1q and loss of 5q as was reported previously^43^.

### Timing of CNVs, ECA and MRCA

Quantification of mutation densities at copy number gains was performed using the R package “NBevolution” v0.0.0.9000, which is described in detail in corresponding study^18^. Briefly, we counted clonal mutations separately on each autosome, stratified by copy number state using the function *count.clonal.mutations()* with max.CN=4, excluding chromosomal segments with length <10^7^ bp. *count.clonal.mutations()* fits a Binomial mixture model with success probabilities according to the expected mean values of the clonal VAF peaks, which, for an impure sample with tumor cell content *ρ*, are given by

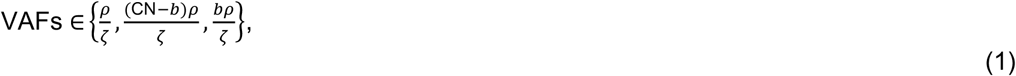

where CN denotes the copy number of a given segment, *b* denotes the number of B alleles on this segment, and

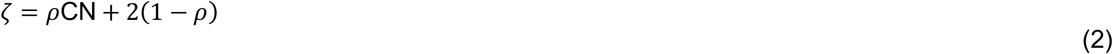

is the average copy number of a given locus in the sample. Mutation densities (SSNVs/bp) at MRCA and ECA, denoted by 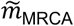 and 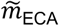 respectively, were computed using the function *MRCA.ECA.quantification()*. In brief, *MRCA.ECA.quantification()* first estimates 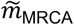 from the number of all clonal mutations and the total size of the analyzed genome, *g* = ∑_*l*_ *g*_*l*_, where the index *l* labels individual segments contributing to the analysis, yielding

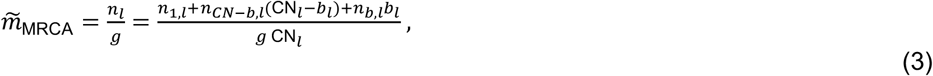

where *n*_*k,l*_ denotes the number of mutations present on *k* copies of the *l*-th segment. Moreover, *MRCA.ECA.quantification()* computes lower and upper 95%-confidence bounds for 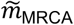 by bootstrapping the genomic segments 1000 times. Next, *MRCA.ECA.quantification()* asks for evidence for an earlier common ancestor in the data. To this end, the function tests for each gained segment with a negative binomial distribution whether amplified clonal mutations agree with the mutation density at MRCA or whether they are significantly less frequent. According to the test result, each segment is either assigned to the MRCA or to an earlier time point. From the latter, *MRCA.ECA.quantification()* computes the mutation densities at the ECA as

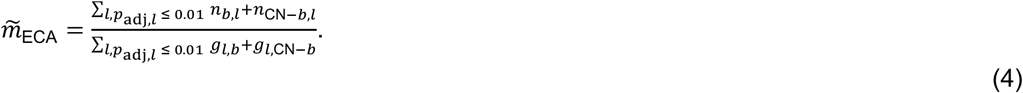

Finally, *MRCA.ECA.quantification()* tests for each contributing segment with a negative binomial distribution whether its mutation density conforms to the ECA and, in analogy to 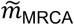, computes lower and upper 95%-confidence bounds by bootstrapping.

Upon timing MRCA and ECA for each sample, we translated mutation densities into weeks post conception (p.c.) by inferring SSNV rates per diploid genome and embryonic day (μλ), using the measured VAF distributions and age at diagnosis as outlined below in section ‘Real-time estimate of cell division rate’. As mutation calling was performed by comparing tumors against a matched blood control, mutation densities correlate with the time post gastrulation (at approximately 2 weeks after conception). Thus, the mutation density per haploid genome, 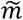, relates to the time p.c. according to 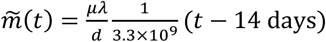.The estimated time of birth was taken as 38 weeks after gastrulation (40 weeks p.c.).

### Timing of SNVs and small indels

We classified single nucleotide variants and small indels as subclonal or clonal based on the number of variant reads, *n*_var_, the number of reference reads, *n*_ref_, tumor purity *ρ* and copy number *k*. Specifically, mutations were classified as subclonal if the probability to sample at most *n*_var_ variant reads out of *n*_var_ + *n*_ref_ total reads according to a binomial distribution with success probability 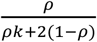. Was smaller than 5%. If a mutation was classified as clonal and fell on a region with *k* = 3, we moreover classified the mutation as early clonal (i.e., acquired prior to the chromosomal gain on the gained chromosome and hence present on two copies) if the probability to sample at most *n*_var_ variant reads out of *n*_var_ + *n*_ref_ total reads was at least 5% according to a binomial distribution with success probability 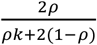.If a mutation was classified as clonal and fell on a region with *k* > 3, we classified the mutation as late clonal (i.e., acquired after the chromosomal gain and hence present on a single copy only) if the probability to sample sample at most *n*_var_ variant reads out of *n*_var_ + *n*_ref_ total reads was smaller than 5% according to a binomial distribution with success probability 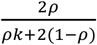, and else as early clonal (acquired prior to the chromosomal gain and hence present on two copies).

### Modeling medulloblastoma initiation

We modeled medulloblastoma initiation and growth with a population-genetics model originally developed for neuroblastoma, as described previously^18^. In brief, the model assumes that disease initiation is driven by 2 consecutive drivers in a transiently expanding tissue of origin, which for G3/4 medulloblastoma is likely the population of differentiating unipolar brush cells (UBCs)^19-21^ . The two driver events are associated with the ECA and the MRCA of the tumor, and spawn, respectively, a pre-malignant and the malignant tumor clone. We assumed that both drivers occur with small probabilities μ_1_ and μ_2_ during cell divisions, and confer a selective advantage (*r* and *s*, respectively), that acts by reducing cell differentiation. Moreover, we assumed that UBCs acquire on average μ neutral somatic variants per cell division, which we modeled with a Poisson process. The population of UBCs has been experimentally described from week 9p.c. until the time of birth^19,23^. To capture this trend, we modeled an initial phase of exponential growth at rate λ_1_ – *δ*_1_, λ_1_ > *δ*_1_ until time *T*, where λ_1_ and *δ*_1_ denote the division and differentiation rate, respectively, and a subsequent phase of exponential decline at rate λ_2_ – *δ*_2_, λ_2_ < *δ*_2_.

Following Körber et al.^18,44^, we calculated the probability of the MRCA to occur at time *t* according to

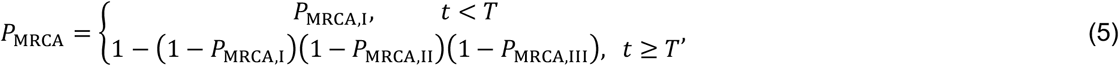

with^40^

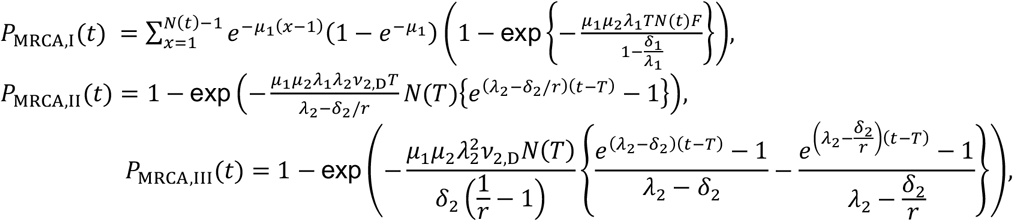

Where 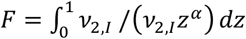 and 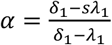. Moreover, we calculated the probability of the ECA to occur at *t*_1_, conditioned on the MRCA occurring at *t*_2_, as described previously^18^

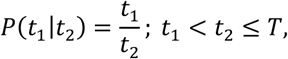

and

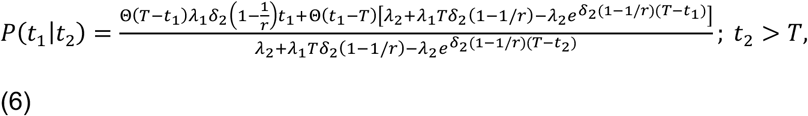

where Θ(·) is the Heavyside step function and 0 ≤ *t*_1_ < *t*_2_.

To estimate the model parameters from the WGS data, we contrasted the probability of acquiring the first and second driver with the measured distribution of SNV densities at ECA and MRCA in G3/4 MBs using Approximate Bayesian Computation with Sequential Monte-Carlo sampling (ABC-SMC) as implemented in pyABC^45^. We used a population size of 1,000 parameter sets and 25 SMC generations or ε ≤ 0.05 as termination criteria. The model fit was performed in analogy to Körber et al.^18,44^ (code and pseudo-code are available on https://github.com/kokonech/mbOncoAberrations). 95% posterior-probability bounds for the model fits were estimated by simulating the model at each sampled parameter set and cutting off 2.5% at each end of the simulated distribution.

### Modeling medulloblastoma growth

We modeled medulloblastoma growth from the MRCA as exponential growth with rate λ_*T*_ – *δ*_*T*_, where λ_*T*_ denotes the division rate and *δ*_*T*_ the loss rate (due to differentiation or death) in the tumor. Assuming that neutral mutations are on average acquired at a rate μλ_*T*_*N*_*T*_(*t*) per haploid genome during tumor growth, the site frequency spectrum of neutral variants at *t*_end_ is, on average^46^,

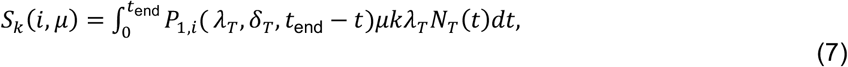

where k is the ploidy, *P*_1,*i*_(λ_*T*_, *δ*_*T*_, *t*_end_ − *t*) is the probability to grow from a single cell to a clone of size *i* within a time span *t*_end_ − *t*, according to a birth-death process (see for example here^47^).

In order to estimate μ and δ_*T*_/λ_*T*_ from the WGS data, we followed the strategy described previously^18^ to compared the cumulative variant allele frequency histogram

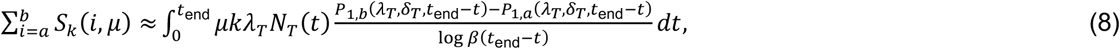

1. between model and data. To this end, we used ABC-SMC^45^ with a population size of 1,000 parameter sets and 25 generations or ε ≤ 0.05 as termination criteria. To learn the dynamics of tumor growth with confidence, we included tumors with well-defined subclonal tails and no evidence for subclonal selection. Tumors were selected (**Extended Data Table 3**) based on visual inspection of the VAF histograms, to remove cases with poor subclonal resolution. In addition, we removed cases with evidence for subclonal selection as suggested by the evolutionary model implemented in Mobster^48^ (setting autosetup = “FAST”), which we ran on autosomes and upon computing pseudo-heterozygous VAFs, 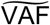, defined as 50% of the mutant sample fraction, SF (hence 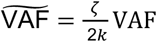, where *k* is the number of alleles carrying the mutation). For the 39 retained G3/4 medulloblastomas, we followed the strategy outlined in Körber et al.^18,44^ to estimate the model parameters from the measured VAF distribution (code and pseudo-code are available on https://github.com/kokonech/mbOncoAberrations ).

### Real-time estimate of cell division rate

From the model fits of medulloblastoma initiation and growth to WGS data, we estimated differentiation/loss rates and mutation rates relative to the rate of cell divisions. To convert these estimates into real-time, we used the age distribution at diagnosis for calibration. In a first step, we estimated the cell division rate of UBCs, λ, from the number of generations between gastrulation and MRCA plus the number of generations between MRCA and diagnosis (*t*_D_), which can be inferred from the mutational burden in the tumor^18^. With λ_*T*_ = *s*λ, this yields

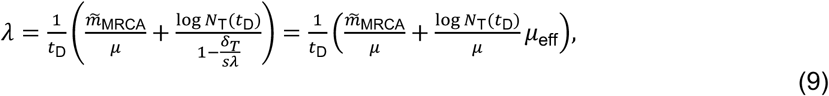

where we used the estimate for μ from the parameter inference for medulloblastoma initiation and the estimate for the effective mutation rate, 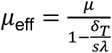, from the parameter inference for medulloblastoma growth. Assuming a tumor mass in the order of a few cubic centimeters and hence *N*_T_(*t*_D_) = 10^9^ cells, and defining *t*_D_ as the age at diagnosis, *A*, plus, on average, 250 days of embryogenesis after gastrulation, we obtained for each tumor (labeled with index *i*) an estimate for the division rate with mean,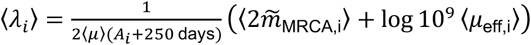, and standard deviation,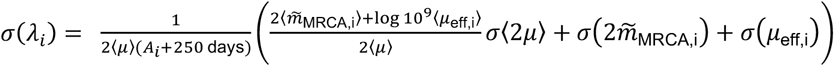, in actual time (where the factor 2 accounts for the fact that μ And 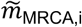 measure the mutation rate and the mutation density, respectively, per haploid genome).

Finally, we computed the mutation rate per day during tumor initiation, by computing μλ_T,i_, with associated uncertainty μΔλ_T,i_ + λ_T,i_σ(μ), which relates molecular clock to real time. For this purpose, we average across the inferences from all tumors that went into the analysis.

## Data availability

The DNA whole genome sequencing mutation results were integrated from the corresponding medulloblastoma molecular landscape study^3^ deposited at European Genome-Phenome Archive under accession number EGAS00001001953. Single nuclei RNA and ATAC data and quality controls published in Joshi *et al bioRxiv* and are available at GEO database under the accession numbers GSE253557 and GSE253573 accordingly. All raw images and processed data after cell segmentation from spatial transcriptomics experiments available at GEO database and can be accessed under the accession number GSE252090.

### Source code

All custom Python/R scripts as well as details about external software environment applied during the data analysis are shared via public github repository: https://github.com/kokonech/mbOncoAberrations

## Author Contributions

K.O., P.J. and V.K. performed main data processing and multi-level bioinformatics analysis. A.R. M.B, and J.P.M. contributed with spatial data generation and analysis. P.B.G.S., B.S., J.V., A.W. and L.K. prepared tumor materials. M.S., I.S., T.Y. and H.K. contributed to the project from evolutionary biology perspective. K.S., M.B.J., P.F., B.J., T.M., K.P., C.M.v.T., O.W., K.B., K.J.W., L.N., C.R., U.S., M.M., S.R. and D.T.W.J. provided tumor tissue samples and feedback for the manuscript. A.K., K.R., F.W., S.T., and T.H. contributed to the design of the study. K.O., P.J., V.K., L.M.K. and S.M.P. prepared the figures and wrote the manuscript based on feedback from all authors. L.M.K, and S.M.P. co-led the study.

## Acknowledgments

This project has received funding from the European Research Council (ERC) under the European Union’s Horizon 2020 research and innovation programme (grant agreement no. 819894) and Seventh Framework Programme (FP7-2007-2013) (grant agreement no. 615253).

The INFORM program is financially supported by the German Cancer Research Center (DKFZ), several German health insurance companies, the German Cancer Consortium (DKTK), the German Federal Ministry of Education and Research (BMBF), the German Federal Ministry of Health (BMG), the Ministry of Science, Research and the Arts of the State of Baden-Württemberg (MWK BW); the German Cancer Aid (DKH), the German Childhood Cancer Foundation (DKS), RTL television, the aid organization BILD hilft e.V. (Ein Herz für Kinder) and the generous private donation of the Scheu family. The study was in parts funded by the European Horizon 2020 Programme - ‘iPC - individualizedPaediatricCure’ (Grant nr. 826121) to S.M.P and by The Medulloblastoma Initiative to L.M.K and S.M.P. We also would like to express our sincere thanks to Carsten Maus, Erjia Wang (Genomics and Proteomics Core Facility, DKFZ), Lena Weiser, Gregor Warsow (Omics IT and Data Management Core Facility, DKFZ) for their highly dedicated support in data management and processing, Robert Autry, Gnanaprakash Balasubramanian, Christopher Previti and Rolf Kabbe (Division of Pediatric Neurooncology, DKFZ) for their sincere and dedicated contribution to the bioinformatics analyses.

